# Rates, Ripples, and Responsivity: The Geometry of Echolocation in Water-Foraging Bats

**DOI:** 10.64898/2026.01.10.698675

**Authors:** Ravi Umadi

**Author notes:** **For correspondence:** (RU).

## Abstract

Water-surface foraging is a rare strategy among echolocating bats, requiring precise coordination between sonar emission, echo timing, and flight geometry. Here, I develop a geometric–responsivity framework to explain how bats foraging over water regulate call timing under these constraints. The model links beam projection, flight height, and specular surface reflections to predictable call-rate scaling and near-field interference patterns. Field recordings of *Myotis daubentonii* reveal spatially invariant spectral ripples across microphones, confirming a geometric interference origin rather than receiver-dependent effects. Reanalysis of classic data from *Noctilio leporinus* shows that call rates are inconsistent with continuous prey-distance locking and instead reflect regulation relative to stable surface-related echoes, with prey-centred control emerging only at close range. Together, these results demonstrate that water-foraging bats structure call timing using environmental reference echoes, providing a mechanistic explanation for the acoustic constraints that shape this specialised foraging niche.

## Introduction

The taxonomic order *Chiroptera*, encompassing all bat species, constitutes an ecologically diverse and behaviourally intriguing group of mammals, comprising over 1,300 species (***Wilson and Mittermeier, 2019***). Most of these species use echolocation as their primary sensory mechanism for orientation, navigation, and prey detection and pursuit. A unique and highly specialised behavioural trait is foraging over water bodies for insects and aquatic animals. Although lakes, rivers, and ponds often support high insect densities (***Speight et al., 2008***; ***Baxter et al., 2005***; ***Nakano and Murakami, 2001***) and provide relatively unobstructed flight corridors, only a small, taxonomically restricted subset of species consistently exploits prey at or just above the water surface. In the Palaearctic region, this behavior is characteristic of trawling bats within the genus *Myotis*, particularly *Myotis daubentonii* (Kuhl, 1817) (***Bogdanowicz, 1994***), *M. dasycneme* (Boie, 1825), and *M. capaccinii* (Bonaparte, 1837) (***Jones and Rayner, 1988***; ***Medard and Guibert, 1990***; ***Britton et al., 1997***; ***Rydell et al., 1999***; ***Siemers et al., 2001***; ***Aihartza et al., 2008***), as well as the fish-eating *Myotis vivesi* (Menegaux, 1901) (***Blood and Clark, 1998***; ***Burt, 1932***; ***Altenbach, 1989***).

Outside Europe, analogous strategies have evolved independently. For example, water-surface foraging has been reported in Asia–Pacific species such as *M. adversus* (Horsfield, 1824) (***Jones and Rayner, 1991***), and most prominently in the Neotropical fishing bats *Noctilio leporinus* (Linnaeus., 1758) (***Hood and Jones, 1984***) and *N. albiventris* (Desmarest, 1818), which capture both aquatic insects and small fish (***Suthers and Fattu, 1973***; ***Hartley et al., 1989***; ***Schnitzler et al., 1994***). The rarity of water-surface foraging across more than 1,300 extant bat species (***Wilson and Mittermeier, 2019***) likely reflects not prey limitation, but rather the stringent sensory, biomechanical, and acoustic constraints imposed by the physical properties of the environment (***Griffin, 1958***; ***Fenton, 1990***; ***Neuweiler, 2003***).

A defining feature of water-surface foraging is that calm water constitutes a strong and coherent acoustic boundary. Unlike vegetation or terrain, a smooth water surface behaves approximately as a specular reflector at ultrasonic frequencies, producing predictable mirror-like reflections with high gain (***Brekhovskikh, 1980***; ***Pierce and Beyer, 1990***; ***Kinsler et al., 2000***). From an acoustic perspective, the surface is not merely an absence of clutter but a dominant and persistent spatial reference. At shallow grazing angles, most of the emitted acoustic energy is reflected away from the bat, creating an unusually low-noise environment in which weak echoes from insects above the surface can be detected reliably. At the same time, the deterministic nature of specular reflection establishes a well-defined two-path propagation geometry in which direct and surface-reflected sound paths coexist.

The acoustic geometry has two immediate consequences. First, the relative delay between direct and reflected paths is tightly linked to bat height, beam orientation, and wavelength. Second, when the two paths overlap in time, their superposition produces periodic amplitude modulations in the frequency domain—commonly referred to as spectral ripples. Such ripple-like structure has been reported repeatedly in recordings of *M. daubentonii* foraging over calm water, and is generally attributed to multipath propagation between bat and surface (Kalko and Schnitzler, 1989a; ***Siemers et al., 2001***). Despite their ubiquity, however, ripple patterns have mainly remained descriptive features: their geometric origin has not been formalised in a predictive framework, nor has their behavioural significance been clearly established.

Importantly, the same boundary physics that produces stable and informative interference under calm conditions becomes destabilising as surface roughness increases. Even modest rippling introduces time-varying scatterers that generate fluctuating echoes and transient ultrasonic noise, degrading prey detectability. Field studies show that *M. daubentonii* strongly prefers calm water and avoids foraging over rippled surfaces, even when insect abundance is higher there (***Rydell et al., 1999***). Water-surface foraging is therefore not opportunistic, but depends on maintaining a narrow range of geometric and acoustic conditions under which the surface acts as a reliable reference rather than a source of clutter.

Maintaining these conditions requires active behavioural regulation. Flight height, beam angle, and call design jointly determine the ensonified space and the location of the dominant reflection point on the surface. Mouth-emitting bats such as vespertilionids and noctilionids can dynamically adjust beam width through changes in mouth gape, thereby steering acoustic energy and modulating both the strength and delay of surface reflections (***Schnitzler et al., 1994***). A wider beam brings the effective reflection point closer to the bat, increasing echo delay and returned energy, whereas a narrower beam shifts reflections farther away and reduces their influence. In this view, spectral ripples are not incidental artefacts, but inevitable physical signatures of a controlled spatial-acoustic configuration.

Interpreting such signatures is complicated by the fact that the joint effects of emission, propagation, and recording geometry shape recorded echolocation signals. Echolocation calls are highly directional, and microphone recordings can be distorted by off-axis reception, distance-dependent filtering, and receiver-specific multipath effects. ***Hartley et al. (1989)*** explicitly addressed this issue by positioning microphones to approximate the signal incident upon the target when studying *N. leporinus* (***Hartley et al., 1989***). More generally, it has been emphasised that single-microphone recordings can misrepresent emitted spectra and amplitudes, and that spatially distributed receivers are often required to disentangle source properties from measurement artefacts (***Ratcliffe and Jakobsen, 2018***). For spectral ripples in particular, it remains unresolved whether they arise at the receiver, as suggested by Kalko and Schnitzler (1989b) and subsequently by ***Ratcliffe and Jakobsen (2018)***, or already close to the source through near-field interaction with the water surface.

A related unresolved issue concerns temporal control during water-foraging approaches. Studies of fishing bats reveal highly structured adjustments of call timing, intensity, and spectral composition as bats approach prey (***Hartley et al., 1989***; ***Schnitzler et al., 1994***). Yet in scenes containing multiple potential acoustic references—most notably the water surface and the prey—it remains unclear which reference governs inter-pulse timing. Call rate could scale with height above the surface, implying surface-centred control, or with bat–prey distance, implying prey-centred control. Alternatively, reference selection may shift dynamically across the various phases of the approach.

Recent theoretical work on biosonar responsivity provides a quantitative framework for addressing this problem (***Umadi and Firzlaff, 2025***). In this framework, inter-pulse interval (IPI) is linked to echo delay *T_a_* via a scaling coefficient *k*_*r*_,

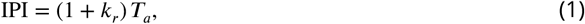

with *T_a_* = 2*d*/*c* for a reference at distance *d* and sound speed *c*. This relationship predicts call rate

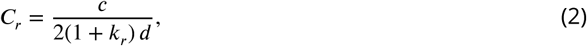

and has been shown to account for call-rate dynamics and timing constraints during aerial prey pursuit (***Umadi and Firzlaff, 2025***). Crucially, because the model makes an explicit distance prediction, it allows the effective acoustic reference to be inferred from measured IPIs: discrepancies between inferred and nominal target distances indicate tracking of an alternative reference.

Here, I integrate geometric modelling, receiver-invariance tests, and responsivity-based timing analysis to clarify the acoustic control principles underlying water-surface foraging. I introduce *Ripple Studio*, a software implementation of a geometric two-path model that predicts spectral ripple spacing as a function of bat height, beam angle, and frequency under specular reflection, incorporating frequency-dependent beam spreading via a circular piston approximation (***Strother and Mogus, 1970***; ***Mogensen and Mohl, 1979***; ***Surlykke et al., 2009***). I then analyse four-channel microphone-array field recordings of *M. daubentonii* to test whether ripple structure is invariant across spatially separated receivers, as expected for a scene-intrinsic phenomenon, or varies systematically with receiver position, as expected for a measurement artefact (Kalko and Schnitzler, 1989b; ***Ratcliffe and Jakobsen, 2018***). Finally, I reanalyse published prey-capture sequences of *N. leporinus* (***Hartley et al., 1989***) within the responsivity framework to determine whether call timing reflects tracking of the water surface or the prey.

By framing spectral ripples and call timing as consequences of controlled spatial-acoustic geometry, this study establishes a basis for interpreting ripple features as behavioural observables rather than incidental artefacts. More broadly, it demonstrates how strong environmental constraints can render biosonar signals invertible, enabling behavioural back-calculation from passive acoustic recordings and providing a principled route for ecological inference in water-foraging bats.

## Material & Methods

### Modelling Spectral Interference in Water-Foraging Bats

To simulate the geometry of acoustic interference resulting from specular surface reflection, I developed a custom MATLAB graphical user interface (GUI). This interface models the two-path propagation of echolocation calls emitted by water-foraging bats and visualises the resultant spectral interference patterns in the received signal.

### Assumptions and Rationale

The model assumes that water-foraging bats emit their echolocation calls downward at a moderate angle toward the water surface, rather than horizontally. This is consistent with behavioural observations of bats such as *M. daubentonii*, which fly low over water while directing their sonar beam slightly ahead of their trajectory. As a result, the primary reflection point lies close to the bat’s position, and the reflected sound returns along a slightly delayed path compared to the direct path to a given interference point (also see Appendix 1).

### Geometrical Construction

The user specifies three parameters via interactive sliders: the bat’s height above the water (ℎ), the beam incidence angle relative to the horizontal (*θ*), and the horizontal distance to the shoreline (*x_shore_*). These inputs define the key spatial locations:

- The bat is positioned at (0, ℎ).
- The reflection point on the water surface is computed as

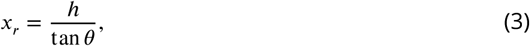

assuming specular reflection and flat geometry.
- The reflected ray is computed by mirroring the incident path across the water surface, yielding an “interference point” located at

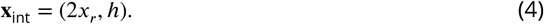 This is the first point at the bat’s height that is reachable both directly and via surface reflection.
- The direct path is simply the Euclidean distance from the bat to this interference point:

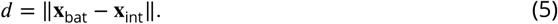
- The reflected path consists of two segments:

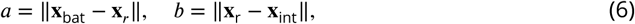

giving a total reflected path length *a* + *b*. The *path difference* is then calculated as:

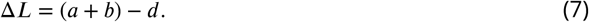

The corresponding acoustic delay is computed using the speed of sound in air (*c* = 343 m/s):

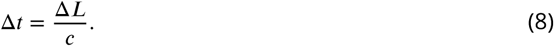

### Spectral Interference Synthesis

To visualise the spectral consequences of two-path interference, a synthetic echolocation call was generated as a frequency-modulated (FM) sweep ranging from 25 to 90 kHz, with a duration of 5 ms and a sampling rate of 192 kHz (refer Umadi (2025f) for call synthesis). The call was duplicated to form a delayed copy, which was shifted in time by a delay Δ*t* corresponding to the path-length difference between the direct and reflected propagation paths. The received signal was modelled as the superposition of the direct and delayed components,

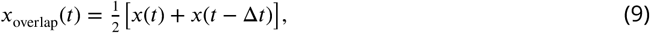

thereby capturing the constructive and destructive interference observed at the receiver when two coherent wavefronts overlap.

A high-resolution spectrogram of the overlapped signal was computed to visualise the resulting interference structure. In the frequency domain, superposition of a signal with a delayed copy produces a periodic comb-like modulation, with spectral minima occurring when the phase difference between the two components equals *π*. Consequently, the fundamental spectral ripple frequency is given by

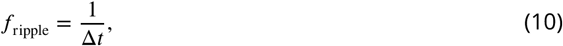

independent of the detailed call structure. This relation directly links the observed ripple spacing to the underlying propagation delay and, consequently, to the geometry of the reflecting surface.

### Beam Width Dynamics and Geometric Overlap Evaluation

To further evaluate the spatial overlap between outgoing calls and returning echoes under realistic foraging conditions, I incorporated frequency-dependent beam width dynamics into the reflection geometry model. The −6 dB beam width at the start and end frequencies of the call was estimated using a circular piston model (***Mogensen and Mohl, 1979***; ***Surlykke et al., 2009***), with the aperture diameter set to the empirical range of mouth gape. This model yields narrower beam widths at higher frequencies, simulating the directional sharpening across the call sweep.

The beam boundaries at both the start and end frequencies were computed by offsetting the main emission angle (*θ*) by half the frequency-specific beam width. Each resulting edge was projected from the emitter using a direction vector defined in polar coordinates. If the upper beam bound exceeded the horizontal plane (i.e., *ϕ* < 0), the projection was clipped to a fixed length, indicating no intersection with the reflective water surface. Otherwise, the beam boundary was extended to the point of water-surface intersection, and a reflected path was rendered symmetrically.

### Contrasting Geometric Model Without Reflection Symmetry

To contrast with the reflection-symmetric model described above, I implemented a second simulation approach based on an earlier geometric interpretation of Kalko and Schnitzler (1989b). This approach assumes that water-foraging bats receive interference patterns resulting from the interaction between direct and surface-reflected paths, but it does not strictly adhere to the law of reflection. Instead, it calculates a triangle formed by the slanting emitted beam and the water surface, intersecting a fixed receiver positioned at a specified horizontal distance (the “shore distance”).

### Assumptions and Geometry

In this model, bats are simulated at various heights ℎ ∈ [0.1, 1.0] m above the water surface, emitting downward-oriented beams at angles *θ* ∈ [5◦, 45◦] below the horizontal. The horizontal distance to the microphone (e.g., the shoreline or an array) is fixed at *x*_mic_ = 5 m.

The geometry proceeds as follows:

- The bat is located at (0, ℎ), and the beam is projected down to the water surface at angle *θ*.
- The reflection point is calculated according to equation (3) assuming the beam strikes the water surface at this horizontal distance.
- The slant path length from the bat to the reflection point is computed as:

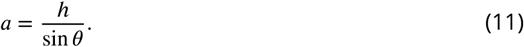
- The second segment, from the reflection point to the receiver (assumed to be at (*x*_mic_, 0)), is:

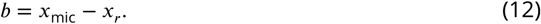 This implicitly assumes a flat, specular water surface but does *not* reflect the beam symmetrically according to the angle of incidence.

### Direct and Indirect Paths

The total indirect path is *a* + *b*, while the direct path is calculated geometrically using the law of cosines:

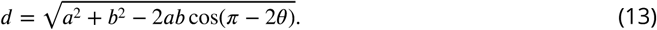

Note that this step introduces an approximate notion of the angle of arrival, but does not model true specular reflection. The path difference is given by equation (7), and the corresponding delay is given by equation (8) with *c* = 343 m/s.

### Spectral Simulation

As in the main model, a virtual echolocation call is generated as a downward FM sweep (25–90 kHz, 5 ms duration), sampled at 192 kHz. This call is delayed according to Δ*t*, and overlapped with the original to simulate the effects of interference, as stated in equation (9).

### Interpretation and Limitations

This model predicts that only a narrow combination of bat height and receiver distance will produce strong spectral interference. The implication is that such interference cues are fragile and geometrically constrained. However, this outcome is a direct consequence of not enforcing the law of reflection. By failing to simulate specular symmetry, the model incorrectly suggests that interference is only observed when the reflected beam happens to reach a distant point (e.g., the microphone) in a rather specific geometry (see Appendix 2).

Nonetheless, this perspective has been influential in the literature (e.g., Kalko and Schnitzler (1989b); ***Ratcliffe and Jakobsen (2018)***), and variants of this model are still commonly referenced. The implementation here serves to highlight the limitations of this approach, and to visually demonstrate the consequences of this simplification when compared to reflection-symmetric models.

### Interactive Simulation of Beam Reflection and Spectral Interference

To support interpretation of the geometric interference model and to facilitate hypothesis testing, all simulations were implemented in an interactive MATLAB application (*Ripple Studio*; figure. 10). The tool instantiates the analytical framework described above and allows real-time exploration of how bat height, beam geometry, and call parameters jointly determine specular reflection, path delay, and the resulting spectral interference patterns.

*Ripple Studio* computes reflection geometry, path-length differences, and overlapped call waveforms for user-defined configurations, and visualises the corresponding time-domain signal, spectrogram, and beam–surface geometry. The simulation is used here as an exploratory and interpretive aid rather than as an independent analytical method.

A detailed description of the interface, parameters, and usage is given in Appendix 3.

### Field Recordings

To validate the predictions from the interference and responsivity models, field recordings were conducted on three consecutive nights along the shoreline of a small lake on the Weihenstephan campus of the Technical University of Munich. Recordings focused on *M. daubentonii* and were performed during the post-sunset activity peak between 20:45 and 21:45.

Acoustic data were collected using the BATSY4-PRO ultrasonic multichannel recording system (Umadi, 2025c), equipped with real-time heterodyne audio output for live monitoring. The DAC output from the BATSY4-PRO was transmitted wirelessly via a Bluetooth A2DP travel adapter to headphones for field guidance. Recordings were triggered using a custom ESP-NOW-based remote controller—an extension developed for the BATSY4-PRO—which allowed operation in either *buffer-dump* mode or *buffer + 10 s record* mode depending on real-time behavioural cues.

A three-dimensional microphone array was deployed and characterised using the Widefield Acoustics Heuristics (WAH) framework (Umadi, 2025f), which provides spatially resolved estimates of localisation uncertainty across the array volume. The highest localisation precision was achieved in the lower region of the array. Accordingly, the array was mounted 2.1 m above the water surface and tilted backwards by 30◦, oriented toward the mid–far field of the lake to maximise detection sensitivity. Each array arm measured 1 m from the centre (see Umadi (2025c) for construction details).

Although localisation algorithms were explored using the multichannel recordings, reliable position estimates could not be obtained for the majority of calls. This limitation arises from a combination of factors intrinsic to water-surface foraging: (i) strong two-path interference patterns between direct and surface-reflected echoes, (ii) rapid fluctuations in call intensity and beam orientation, and (iii) variable relative flight velocity with respect to individual microphones in the array. Together, these effects violate key assumptions underlying conventional time-difference-of-arrival and beamforming approaches, leading to unstable or ambiguous localisation solutions (Umadi, 2025f). Subsequent analyses focused on call timing, spectral structure, and responsivity-based inference, which remain robust under these recording conditions.

### Echolocation Call Detection and Segmentation

Echolocation calls were segmented from each multichannel waveform using the *Biosonar Responsivity Analysis Toolkit* (Umadi, 2025b) in combination with the *Array WAH: Microphone Array Design Tool* (Umadi, 2025a,f). Each detected call was retained only if its peak amplitude exceeded a pre-defined quality threshold (0.0035 relative amplitude), ensuring reliable measurements.

Signals were aligned across channels by cross-correlating each channel with Channel 1 and applying the corresponding lag correction. To minimise onset variability, each channel waveform was further time-shifted so that its maximum peak was centred in the analysis window.

### Quantitative Analysis of Amplitude Modulation and Ripple Metrics

I implemented a custom MATLAB analysis pipeline to characterise both the amplitude modulation (AM) patterns and the ripple interference frequency (*f*_ripple_) on a per-channel basis. For each extracted call:

1. **Envelope extraction:** The analytic signal was obtained using the Hilbert transform, and its magnitude was smoothed with a 4th-order zero-phase Butterworth low-pass filter (cutoff 5 kHz).
2. **Call duration:** The call segment was defined between the 10% and 90% cumulative energy bounds (function percentdur), yielding the duration (Dur*_*ms*_*), onset (*t_*L*_*), offset (*t_*R*_*), RMS amplitude, and peak frequency (*f_*max*_*).
3. **AM frequency (*f*_AM_):** Within the extracted segment, the normalised envelope was analysed using Welch’s power spectral density estimate (window length 512 samples, 50% overlap, 4096-point FFT). The dominant spectral peak in the 0.3–20 kHz band was taken as *f*_AM_. If no stable spectral peak was detected, a fallback autocorrelation method was applied.
4. **Ripple delay and frequency (*τ*, *f*_ripple_):** To quantify interference between direct and reflected paths, I applied three complementary methods implemented in the *Ripple Studio* framework:
  - *Time-domain cross-correlation*: locating the strongest secondary correlation peak within 0.2–2.0 ms lags.
  - *Real cepstrum analysis*: identifying peaks in the cepstrum that correspond to periodicities in the modulation envelope.
  - *Spectral comb analysis*: band-pass filtering the waveform (20–90 kHz), computing its spectrum, and detecting periodicities in the spectral envelope. The spacing of comb peaks corresponds to the ripple frequency.
  A consensus echo delay (*τ*_cons_) was derived as the median of the consistent estimates across methods, and the ripple frequency was calculated as *f*_ripple_ = 1000/*τ*_cons_ (Hz).
5. **Duration inflation:** For calls with clear evidence of overlap (i.e. where at least two independent methods agreed within 0.15 ms), an additional metric was computed as the increase in measured call duration attributable to overlap:

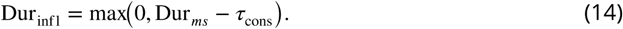

For each file and channel, all metrics (duration, corrected duration, envelope-derived *f*_AM_, ripple frequency *f*_ripple_, peak prominence, and RMS level) were stored in a summary table. Subsequent statistical analyses included Kruskal–Wallis tests for channel effects on each parameter, post-hoc comparisons with Dunn–Šidák correction (***Šidák, 1967***), and effect size estimation (*η*^2^, *ε*^2^). Pairwise correlations between call duration and both *f*_AM_ and *f*_ripple_ were also computed (Pearson’s *r* and Spearman’s *ρ*) to quantify associations between call structure and interference patterns.

### Reanalysis of Hartley et al. (1989) Call–Distance Data

To evaluate whether echolocation call timing in trawling bats reflects continuous prey-distance tracking or responsiveness to a nearer acoustic reference, I reanalysed behavioural data reported by ***Hartley et al. (1989)*** within the responsivity framework developed here. Figures 14 and 15 from the original publication were digitised using *PlotDigitizer v3.3.9* (por, 2026), yielding paired measurements of reported target distance (bat height above water) and either call rate (Hz) or inter-pulse interval (IPI; ms) for two individual bats.

Call-rate and IPI data were digitised from separate figures in the original study and treated as independent measurements, rather than algebraic inverses, which reduced sensitivity to digitisation bias and allowed for internal consistency checks. All calculations assumed a sound speed of *c* = 343 m s_−1_.

### Responsivity model

Within the responsivity framework, call timing is governed by a scaled echo delay, such that the inter-pulse interval (IPI) increases proportionally with the acoustic two-way travel time (equation 1). The corresponding prediction for call rate follows directly from this relationship (equation 2), yielding a hyperbolic dependence of call rate on the reference distance governing timing control.

### Estimation of *k*_*r*_ from call-rate data

Rather than fitting *k*_*r*_ to inter-pulse intervals as a function of reported prey distance—which implicitly assumes continuous prey-distance locking—I estimated *k_*r*_* directly from call-rate data. This approach isolates the behavioural regime in which the timing control law is most clearly expressed and avoids imposing prey tracking *a priori*.

For each bat, a near-target (“terminal”) subset was defined as the closest 20% of reported distances. Within this subset, the 95th percentile of observed call rate was taken as a robust estimate of the maximum achieved call rate (C_r_ ^∗^). A representative distance *d*^∗^ was defined as the median of the same subset, reducing sensitivity to outliers and digitisation noise. An anchored estimate of responsivity was then computed as

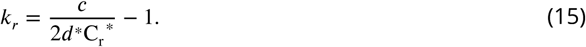

To assess sensitivity to digitisation uncertainty, bootstrap resampling (***Efron, 1979***) was performed on the terminal subset with additive Gaussian noise applied to both distance (*σ_*d*_* = 0.02 m) and call rate (*σ*_C_r__ = 3 Hz). Confidence intervals were derived from the resulting bootstrap distribution of *k_*r*_*.

### Cross-validation of call-rate and IPI datasets

Because call-rate and IPI measurements were digitised independently, additional cross-validation was performed to verify that inferred cue-distance patterns were not artefacts of missing or unevenly sampled data. For each bat, the union of reported distances across both datasets was constructed. Where a call-rate measurement lacked a corresponding IPI value at the same distance, the IPI was interpolated from the nearest neighbouring measurements within a conservative tolerance (±0.08 m); the same procedure was applied symmetrically to fill missing call-rate values for IPI-only distances.

This procedure ensured that both plots were evaluated over a common distance support while preserving the independence of the original measurements. In practice, nearly all data points could be cross-supported in this manner (see Results), indicating that the observed deviations between inferred cue distance and reported target distance do not arise from incomplete digitisation but reflect structure intrinsic to the original dataset.

### Inversion of IPI to infer effective cue distance

Using the anchored *k_*r*_* value obtained from call-rate data, each independently digitised IPI measurement was inverted to estimate the distance of the acoustic cue governing call timing:

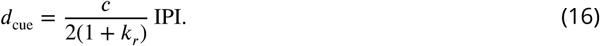

Here, *d*_cue_ represents the inferred *one-way* distance to the reference echo that sets the inter-pulse interval. These inferred cue distances were compared directly with the reported bat–target distances. Systematic deviations below the unity line (*d*_cue_ *< d*_reported_) indicate that call timing was governed by echoes originating substantially closer than the prey itself.

All analyses and visualisations were performed in MATLAB R2023b using custom scripts developed for this study.

## Results

### Cross-Channel Consistency of Modulation Patterns

To assess whether amplitude modulation (AM) patterns produced by near-field interference were preserved across spatially separated microphones, I conducted a cross-channel analysis on the five strongest calls (figure 5). For each call, signals from all four channels were aligned to the reference channel using cross-correlation and subsequently centred on the global amplitude peak. This procedure ensured that comparisons were made on temporally aligned waveforms and envelopes.

The aligned spectra and spectrograms confirmed that the dominant frequency sweep structure was consistently captured across channels, while the raw waveforms showed minor amplitude differences attributable to spatial geometry. Crucially, the normalised AM envelopes revealed a striking similarity in their temporal trajectories, with overlapping rise–fall dynamics across all four channels.

Quantitative comparisons further substantiated this observation (table 1). Pearson correlation coefficients between channel pairs were typically very high (*r >* 0.75, many exceeding *r* = 0.90), while RMSE values were close to zero (∼ 10_−10_). Normalised cross-correlation peaks approached unity with minimal lags (generally within ±0.5 ms), indicating strong temporal alignment of modulation envelopes. Dynamic Time Warping (DTW) (***Sakoe and Chiba, 1978***; ***Müller, 2007***) distances also remained uniformly low (*<* 10^−6^), demonstrating that little to no non-linear alignment was required to achieve close similarity.

**Table 1.** Pairwise comparison of amplitude modulation (AM) envelopes extracted from all four microphone channels across five different recordings of a water-foraging bout reveals a striking consistency in signal structure. High Pearson correlation coefficients (typically *>* 0.75), low RMSE values (∼ 10^−10^), and prominent cross-correlation peaks with minimal lag confirm strong linear and temporal similarity across spatially separated microphones. Crucially, the Dynamic Time Warping (DTW) distances remain uniformly low (∼ 10^−7^), indicating that only minimal non-linear alignment was needed to match envelopes across channels. Together, these metrics provide robust evidence for spatial invariance in the modulation patterns, supporting the hypothesis that the observed spectral ripples are not caused by interference at the receiver, but rather arise from near-field reflections interfering with the emitted call itself—consistent with the law of reflection.

| File | Ch. | Ch. | Pearson Corr. | RMSE | Max XCorr. | Lag (ms) | DTW |
| --- | --- | --- | --- | --- | --- | --- | --- |
| 1 | 1 | 2 | 0.948 | 3.13e-10 | 0.965 | -0.0156 | 2.3e-07 |
| 1 | 1 | 3 | 0.819 | 1.95e-10 | 0.877 | 0.365 | 7.51e-08 |
| 1 | 1 | 4 | 0.795 | 2.02e-10 | 0.862 | -0.198 | 1.12e-07 |
| 1 | 2 | 3 | 0.819 | 3.88e-10 | 0.872 | 0.0625 | 2.53e-07 |
| 1 | 2 | 4 | 0.804 | 4.2e-10 | 0.874 | 0.161 | 3.1e-07 |
| 1 | 3 | 4 | 0.881 | 1.53e-10 | 0.937 | 0.417 | 8.68e-08 |
| 2 | 1 | 2 | 0.924 | 3.65e-10 | 0.972 | -0.224 | 3e-07 |
| 2 | 1 | 3 | 0.852 | 2.32e-10 | 0.897 | -0.224 | 1.2e-07 |
| 2 | 1 | 4 | 0.842 | 1.75e-10 | 0.895 | -0.25 | 7.02e-08 |
| 2 | 2 | 3 | 0.857 | 3.12e-10 | 0.895 | 0.411 | 2.02e-07 |
| 2 | 2 | 4 | 0.852 | 3.54e-10 | 0.903 | -0.229 | 2.52e-07 |
| 2 | 3 | 4 | 0.872 | 2e-10 | 0.948 | -0.323 | 1.1e-07 |
| 3 | 1 | 2 | 0.867 | 4.51e-10 | 0.959 | 0.224 | 3.55e-07 |
| 3 | 1 | 3 | 0.798 | 3.04e-10 | 0.873 | -0.266 | 1.99e-07 |
| 3 | 1 | 4 | 0.79 | 2.04e-10 | 0.862 | 0.26 | 7.08e-08 |
| 3 | 2 | 3 | 0.797 | 3.86e-10 | 0.876 | -0.0677 | 1.88e-07 |
| 3 | 2 | 4 | 0.813 | 4.24e-10 | 0.866 | 0.0417 | 2.83e-07 |
| 3 | 3 | 4 | 0.865 | 2.4e-10 | 0.903 | 0.00521 | 1.57e-07 |
| 4 | 1 | 2 | 0.94 | 2.93e-10 | 0.964 | -0.0312 | 2.7e-07 |
| 4 | 1 | 3 | 0.767 | 2.39e-10 | 0.877 | -0.161 | 1.32e-07 |
| 4 | 1 | 4 | 0.743 | 2.65e-10 | 0.853 | -0.245 | 9.97e-08 |
| 4 | 2 | 3 | 0.798 | 4.3e-10 | 0.868 | -0.0833 | 3.87e-07 |
| 4 | 2 | 4 | 0.751 | 4.04e-10 | 0.835 | -0.172 | 2.86e-07 |
| 4 | 3 | 4 | 0.888 | 1.89e-10 | 0.937 | 0.0677 | 1.26e-07 |
| 5 | 1 | 2 | 0.804 | 4e-10 | 0.944 | -0.661 | 2.55e-07 |
| 5 | 1 | 3 | 0.9 | 2.05e-10 | 0.929 | 0 | 1.62e-07 |
| 5 | 1 | 4 | 0.836 | 2.58e-10 | 0.892 | -0.193 | 9.7e-08 |
| 5 | 2 | 3 | 0.779 | 4.81e-10 | 0.929 | 0.693 | 3.73e-07 |
| 5 | 2 | 4 | 0.802 | 3.79e-10 | 0.895 | 0.464 | 2.28e-07 |
| 5 | 3 | 4 | 0.871 | 2.87e-10 | 0.941 | -0.406 | 2.32e-07 |

To quantify whether amplitude modulation (AM) and ripple measures were preserved across different microphone channels of the array, I analysed all validated (*N* = 207 × 4) *M. daubentonii* calls on a per–channel basis (figure 6). Call durations varied between channels, with Kruskal–Wallis tests indicating highly significant differences (Duration: *p* → 0, *η*^2^ = 0.155). After correcting for estimated duration inflation caused by multipath interference, differences remained but were slightly reduced (Corrected Duration: *p* → 0, *η*^2^ = 0.138). In contrast, channel effects on AM frequency were small but statistically detectable (*p* =→ 0, *η*^2^ = 0.030), and ripple frequency showed significant differences across channels (*p* =→ 0, *η*^2^ = 0.071), although the effect sizes were modest compared with those for duration. Correlation analyses revealed that call duration was negatively associated with *f_*AM*_* (Spearman *ρ* = −0.39, *p* → 0) and positively associated with *f_*ripple*_* (Spearman *ρ* = 0.31, *p* → 0). These results indicate that while amplitude modulation and ripple frequency patterns are broadly consistent across channels, the apparent call durations vary systematically with sensor position, likely reflecting geometric effects of call–echo interference rather than intrinsic signal changes.

Together, these results confirm that the interference-driven modulation patterns are spatially invariant across the array. The preservation of AM structure across channels supports the interpretation that spectral ripples arise from near-field reflections close to the bat, rather than from interference at the receiver. This provides strong experimental validation of the interference model predictions (Methods) and the reflection-compliant framework developed in *RippleStudio*.

### Field Call rates and Responsivity-Inferred Distance

Analysis of 54 field-recorded call sequences (568 validated calls; figure 7) revealed that call rates during water-foraging clustered tightly around low to moderate values, with a mean of 13.5 Hz and a median of 11.6 Hz (range: 4.2–113.4 Hz). These values are highly consistent with classic field measurements for *Myotis daubentonii*: Kalko and Schnitzler (1989a) reported an average inter-pulse interval of 66.7 ± 14.4 ms, corresponding to call rates of approximately 12.3–19.1 Hz (mean ∼15 Hz). Thus, the dominant calling regime observed in the present dataset closely matches established species-typical foraging behaviour.

Applying the responsivity inversion with *k_*r*_* = 5 yielded inferred effective target distances centred around 2–3 m (mean: 2.48 m; median: 2.45 m), with occasional excursions to much shorter ranges associated with transient increases in call rate. Importantly, these inferred distances arise directly from call timing alone, without assuming explicit localisation or prey locking. Together, the call-rate statistics and responsivity-based distance estimates indicate that most calls were produced while bats operated within a stable, mid-range reference regime, punctuated by brief high-rate episodes likely corresponding to short-lived near-field engagements.

### Predicted Iso-Ripple Contours from the Interference Model

I further explored how geometric factors might give rise to the observed channel-specific differences in duration by simulating ripple frequencies across a dense grid of bat heights (0.1–1.5 m) and beam angles (5–60°) using the *Ripple Studio* interference model (figure 8). Iso-frequency contours (1–5 kHz, 1 kHz spacing) reveal extended regions of height–angle combinations that produce nearly identical *f_*ripple*_* values. These degeneracies demonstrate that multiple spatial configurations can generate the same ripple frequency.

### Reanalysis of Hartley et al. (1989)

Figure 9 presents a reanalysis of the call-rate–distance and inter-pulse interval (IPI) data reported by ***Hartley et al. (1989)*** for two *Noctilio leporinus* individuals. Call-rate measurements were used to estimate the responsivity constant *k_*r*_*, which was subsequently applied to invert independently digitised IPI measurements to infer the effective acoustic distance governing call timing.

Anchoring *k_*r*_* to the terminal high-rate regime—defined as the closest 20% of reported distances and the 95th percentile of call rate within that subset—yielded consistent responsivity estimates across individuals. For Bat 1, the anchored estimate was *k_*r*_* = 3.58 (95% CI: 3.15–4.02), while Bat 2 yielded *k_*r*_* = 3.73 (95% CI: 3.18–4.02). Bootstrap resampling indicated that these estimates were robust to digitisation uncertainty. These values are substantially lower than those implied by strict prey-distance locking, given the reported flight heights and observed maximum call rates.

Because call-rate and IPI measurements were digitised from separate figures, additional cross-validation was performed to ensure that missing data points did not bias the comparison. Using a conservative distance-based interpolation to construct the union of both datasets, nearly all call-rate distances could be supported by corresponding IPI values and vice versa (Bat 1: 116/117 call-rate points; 104/106 IPI points; Bat 2: 143/143 call-rate points; 118/120 IPI points). This confirms that the observed patterns do not arise from incomplete digitisation but reflect a structure intrinsic to the original data.

When the call-rate–derived *k_*r*_* (figure 9a) values were used to invert the independently digitised IPI measurements, the inferred cue distances were systematically compressed relative to the reported bat–prey distances (Fig. 9b). Across both bats, the median inferred distance was approximately 42–44% of the reported target distance (Bat 1: median = 0.44, mean = 0.55; Bat 2: median = 0.42, mean = 0.47). All points lay well below the unity line for most of the approach, with convergence toward equality occurring only at reported distances of approximately 0.5 m.

These results demonstrate that the call rates and IPIs reported by ***Hartley et al. (1989)*** are inconsistent with continuous prey-distance tracking throughout the approach. Instead, they are most parsimoniously explained by bats regulating call timing relative to a nearer acoustic reference for much of the flight, with prey-centred control emerging only late in the capture sequence.

Simulation of Spectral Interference with ***Ripple Studio***

All geometric and acoustic simulations presented in this study are implemented in *Ripple Studio*, an interactive application developed to support hypothesis testing and experimental design (figure 10). The app integrates the geometric interference model with responsivity-based timing relationships, allowing users to explore how bat height, beam angle, call structure, and surface geometry jointly determine spectral ripple patterns and effective call rates.

*Ripple Studio* functions as a computational framework that packages the study’s analytical predictions into an exploratory tool. By enabling real-time manipulation of geometric and acoustic parameters, it facilitates evaluation of ensonification strategies, assessment of parameter degeneracies (e.g. iso-ripple conditions), and generation of testable predictions for controlled laboratory or field experiments.

In this way, *Ripple Studio* provides a transparent link between theory and observation, and establishes a reproducible platform for probing how environmental geometry constrains biosonar control in water-foraging bats.

## Discussion

Water-surface foraging poses a fundamentally different control problem from aerial hawking, yet the question of *what acoustic reference governs call timing* in this context has remained largely unexamined. In contrast to aerial foraging environments—where trees, ground, buildings, or clutter provide abundant and spatially persistent reference echoes—open water offers few stable reflectors, and most acoustic energy is specularly reflected away from the bat. Despite this, several species of bats routinely forage over water far from shorelines, indicating that call timing must be stabilised by mechanisms other than continuous prey-distance tracking or fixed environmental landmarks. In this study, I integrate geometric modelling, spectral interference analysis, and the responsivity framework to reinterpret how bats structure ensonification, select temporal anchors, and extract prey-relevant cues during water-surface foraging. This perspective shifts the focus from prey-centred distance tracking to scene-level organisation, in which behavioural control, beam geometry, and echo predictability jointly determine foraging performance.

### Geometric Origin of Spectral Ripples in Water-Surface Foraging

Spectral ripple structure is a long-recognised characteristic of echolocation calls recorded from *M. daubentonii* and related species flying low over water, yet its mechanistic origin has remained contentious. Early work by Kalko and Schnitzler (1989a) demonstrated that regularly spaced spectral minima in frequency-modulated (FM) calls could be predicted by a two-wave-front interference model, in which a direct call recorded at the microphone interferes with a delayed copy reflected from the water surface. Under this interpretation, ripple spacing is determined by the delay Δ*t* between the two wave fronts, such that minima occur at frequencies *f_n_* = *n*∕Δ*t* While this model successfully reproduces observed ripple spacing under specific recording geometries, it implicitly attributes ripple formation to receiver-side interference between the emitted signal and its surface-reflected copy.

Subsequent methodological critiques have highlighted a key implication of this interpretation ***Ratcliffe and Jakobsen (2018)*** emphasised that microphone recordings of bat echolocation are highly sensitive to recording geometry, beam directionality, and spatial position relative to the bat and reflecting surfaces. Apparent spectral structure in recorded signals may therefore arise from measurement artefacts rather than from features present in the acoustic scene as perceived by the bat. In this context, the receiver-side interference explanation of spectral ripples raises an important question: if ripple structure arises primarily from interference at the microphone, should it not vary systematically with receiver position in a spatially distributed array?

The present study directly addresses this question. Using simultaneous recordings from multiple spatially separated microphones, I demonstrate that the ripple structure remains remarkably consistent across receivers, despite substantial differences in microphone position and angle relative to the bat (figure 5). This spatial invariance is incompatible with a purely receiver-dependent interference mechanism, which would predict receiver-specific ripple spacing and depth. Instead, the data indicate that ripple structure is intrinsic to the acoustic scene itself and therefore shared across receivers. This finding resolves a long-standing ambiguity by demonstrating that spectral ripples are not primarily a measurement artefact, but arise from stable physical interactions between the emitted call and the water surface.

The geometric modelling presented here (figure 3) provides a mechanistic explanation for this invariance. By explicitly modelling specular reflection at the water surface and the near-field interaction between outgoing sound energy and its immediate surface reflection, the model shows that interference can occur effectively at the source or within the propagating sound field before the signal diverges toward spatially separated receivers (figure 1). Under this configuration, ripple spacing is governed by bat height, emission angle, and call frequency, rather than by microphone position. Importantly, the model reproduces both the ripple spacing and its stability across receivers observed in the field data (figure 4), providing a forward, predictive account of ripple formation.

**Figure 1.**
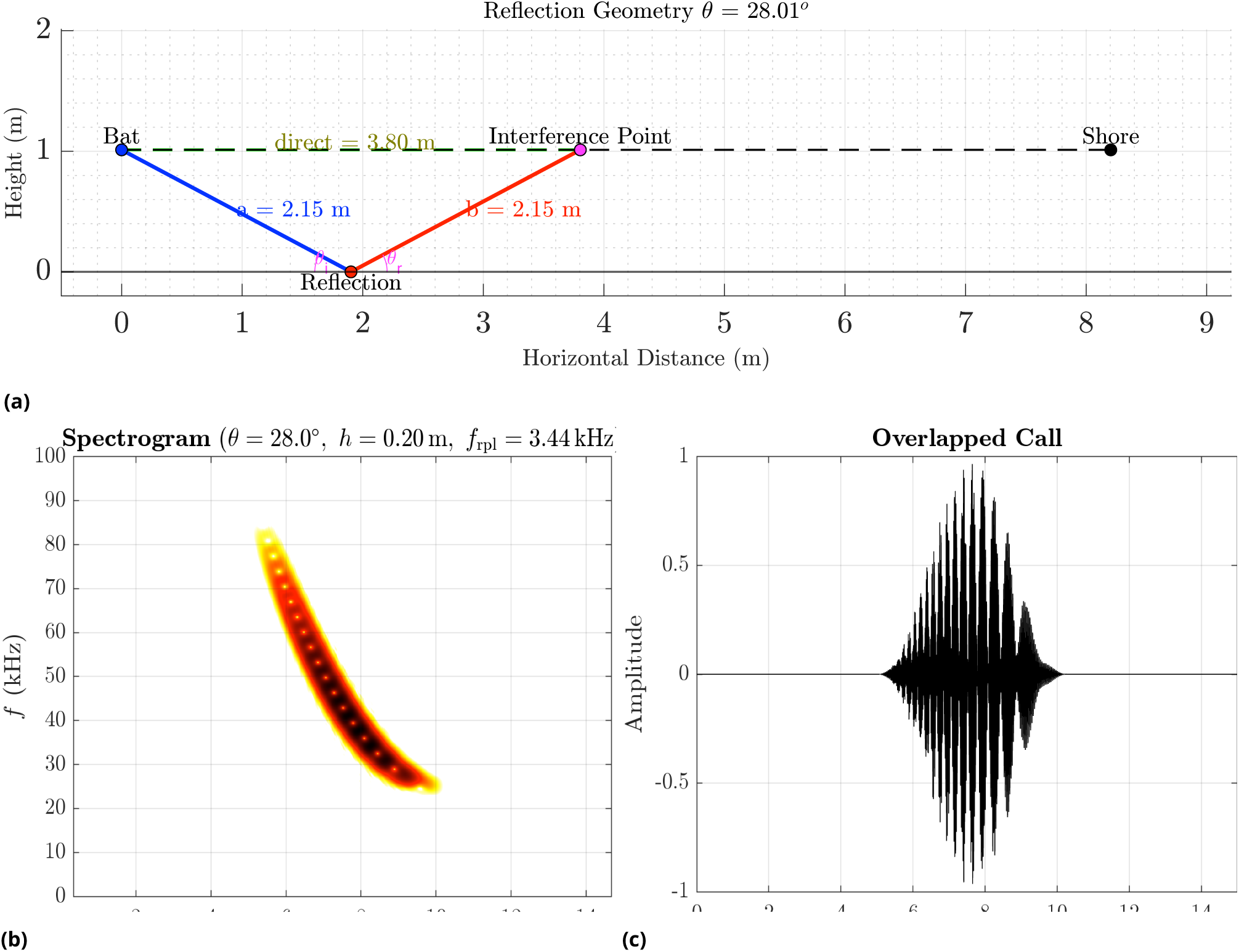
Geometric model and synthetic reconstruction of spectral ripples from surface interference. (a) Illustration of the reflection geometry used in the ripple model: a direct path from the bat to the interference point (dashed), and an indirect path comprising a specular water-surface reflection (blue and red segments). For the shown configuration (reflection geometry *θ* = 28.01◦), the indirect path length exceeds the direct path, producing a fixed path-length (and delay) offset that generates interference. (b) Simulated spectrogram of the resulting overlapped FM call for the same geometry (example: *θ* ≈ 28◦, *ℎ* = 0.20 m), showing the characteristic ripple pattern with ripple spacing *f*_ripple_ = 3.44 kHz. (c) Time-domain waveform of the overlapped call (direct + surface-reflected components) used to generate the spectrogram in (b).

**Figure 2.**
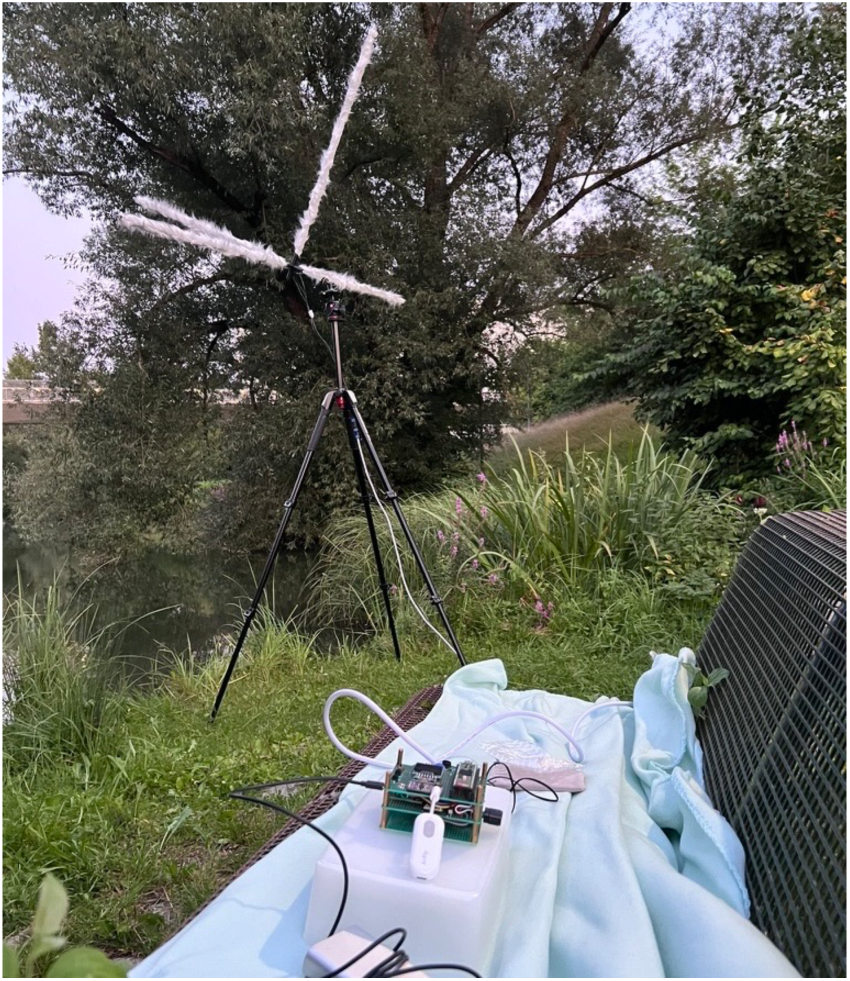
The 3D array was deployed on the shore of the TUM campus lake, mounted on a standard tripod with the top microphone at a height of 2.9 m and the whole array tilted backwards by 30◦ from vertical. The recordings were remotely triggered using an ESP_NOW addon module, and the bat activity was monitored via Bluetooth earphones connected to a commercially available A2DP transmitter plugged into the DAC output of the BATSY4-Pro.

**Figure 3.**
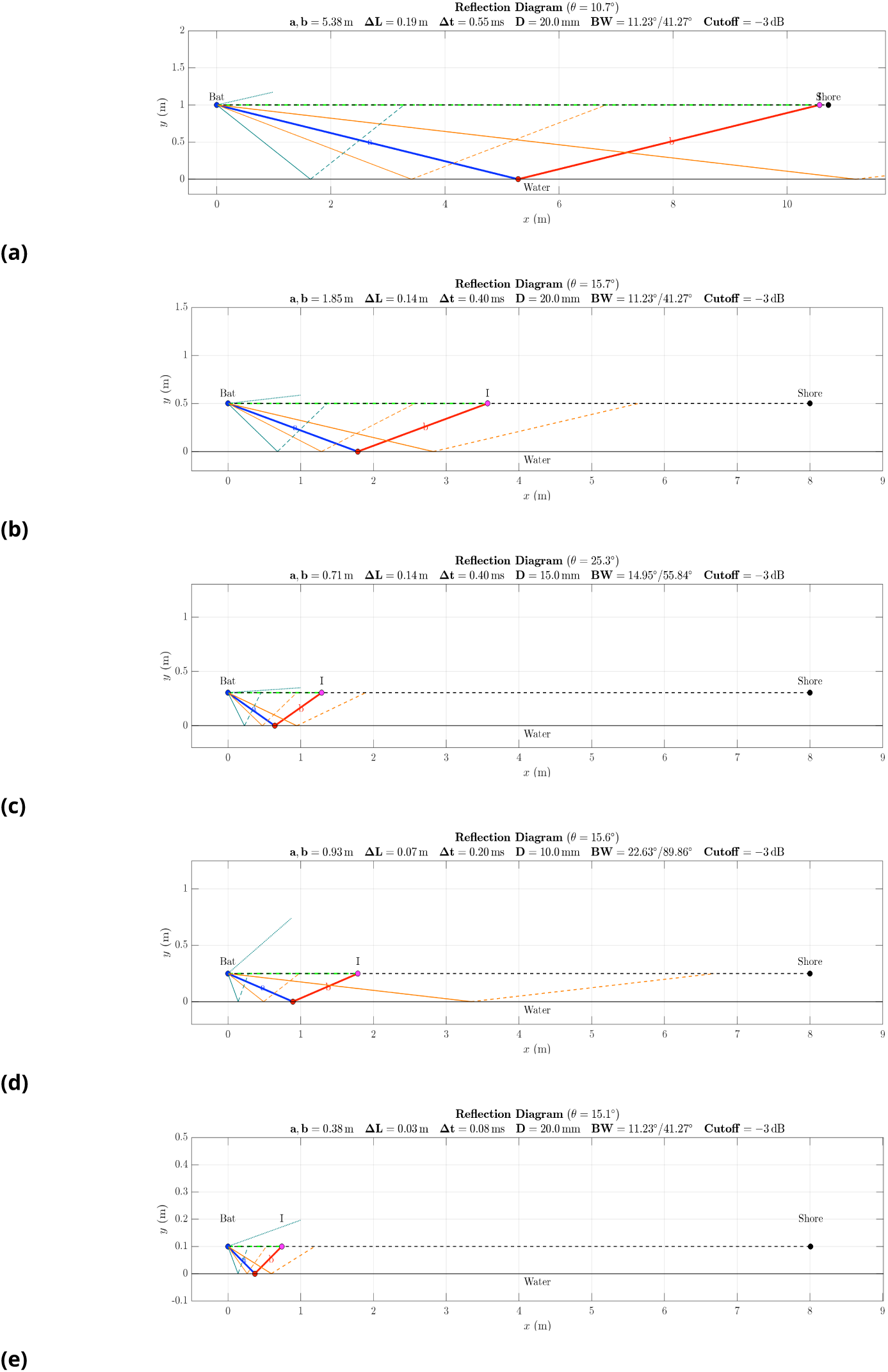
Geometric acoustic analysis showing how combinations of bat height and beam angle create the path differences responsible for spectral ripples in the calls of water-foraging bats such as *M. daubentonii*. The reflection diagrams illustrate how energy emitted at higher frequencies—more tightly focused and occurring earlier in the FM downsweep—projects at shallower angles than the later, lower-frequency energy, at the lower beam boundary, giving rise to interference between direct and reflected paths in the near field. Beam boundaries are drawn at a chosen amplitude drop (–3 dB), but acoustic radiation extends beyond this cutoff, such that reflections and ripples can occur even at considerable heights (e.g. 1 m above the water). When the upper beam boundary crosses the horizontal line, it is drawn as a dotted line of unit length. Different geometries can produce similar ripple patterns, as demonstrated by the spectral features in Figure 8, highlighting that the observed ripples are a robust outcome of beam–surface interaction. Using features of the *Ripple Studio* app, one can observe that ripples emerge not only below 0.3 m bat height, but also at 1 m, with the strongest variations in ripple frequency intervals occurring as the bat descends and beam orientation shifts. In the diagrams, the blue and red lines represent the centre of the beam, oriented at angle *θ*, with arms *a*and *b* forming the reflected path compared against the direct path (green), which intersects the reflection at *I* with path difference Δ*L* and acoustic delay Δ*t*. The aperture *D* defines the beam widths (*BW*) at the start and end frequencies, measured at the chosen cutoff, based on the standard piston model of acoustic radiation. Notably, recordings beyond the point of interference—typically close to the bat—are spatially invariant, such that microphone position does not affect the observed ripple patterns.

**Figure 4.**
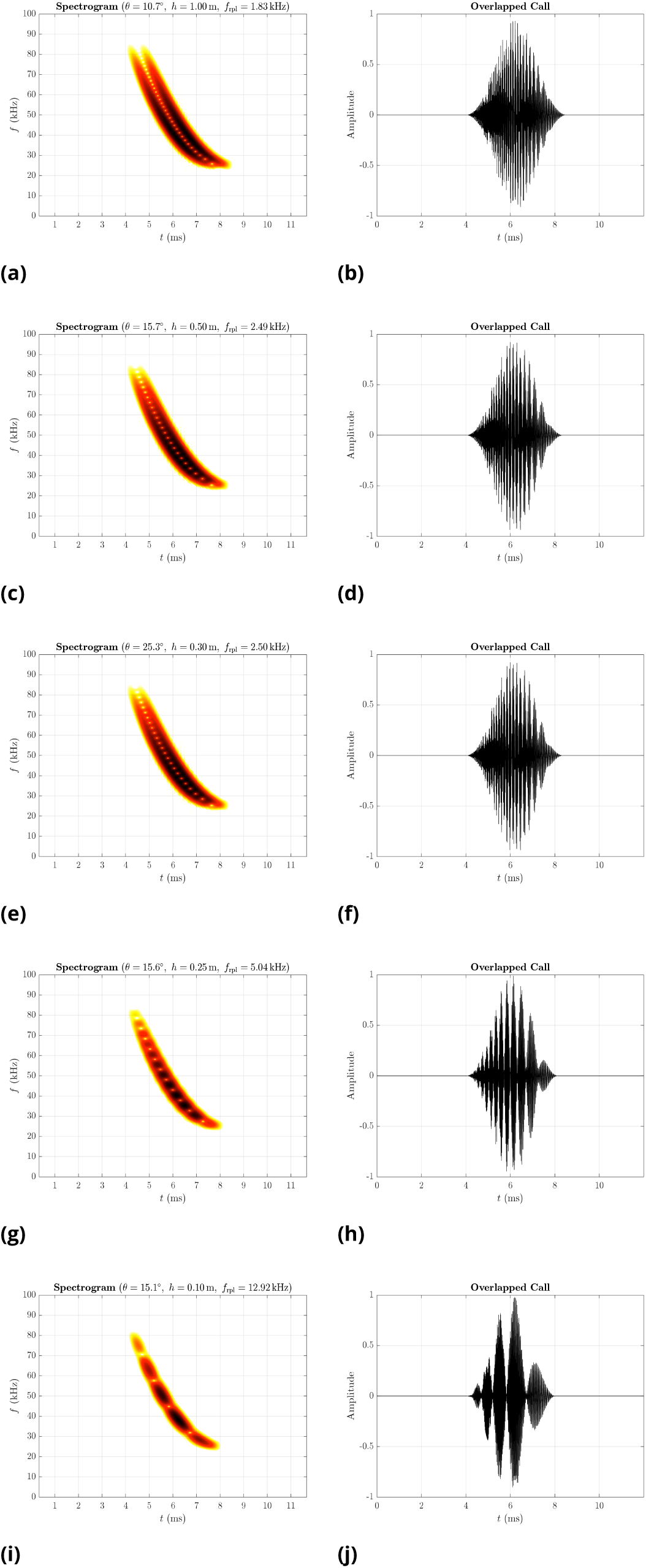
Synthetic echolocation calls illustrating spectral ripples generated by interference between direct and reflected paths. Each row displays the spectrogram (left) and corresponding overlapped waveform (right) for a single geometry, aligning with the reflection diagrams in figure 3. Variations in path difference Δ*L* produce systematic changes in ripple spacing and effective amplitude modulation frequency, whereas overall call duration does not alter the ripple pattern.

This interpretation reconciles earlier empirical observations with modern methodological constraints. Kalko and Schnitzler (1989b) two-wave-front formulation correctly identified the role of surface-reflected energy in shaping ripple structure, but their receiver-centred framing was perhaps a pragmatic simplification dictated by available recording techniques at the time. The present results extend their insight by showing that the relevant interference does not require the microphone to participate in the interference geometry.

By resolving the origin of spectral ripples as a consequence of predictable geometric interactions between the bat’s emission and the water surface, this study establishes ripples as informative acoustic features rather than incidental artefacts. Because ripple spacing depends systematically on bat height and emission geometry, ripple structure encodes information about the spatial configuration of the foraging bat. This provides a physical basis for the backward interpretation developed here, in which spectral features in field recordings can be used to infer aspects of flight height and ensonification strategy. More broadly, the results demonstrate how strong environmental constraints—here, specular reflection from a calm water surface—can imprint stable, interpretable structure onto biosonar signals, thereby enabling ecological inference from passive acoustic data.

### Ensonification, Reference Echoes, and Prey Detection over Water

A central paradox of water-surface foraging is that successful prey detection must occur within a closed-loop echolocation regime that is temporally governed by a stable acoustic reference. Under the responsivity framework, call timing is adjusted relative to the delay of a dominant reference echo, ensuring temporal predictability and sensorimotor stability. For bats foraging over water, this reference is most plausibly provided by the specularly reflecting water surface itself. Prey detection, however, requires that echoes from insects—whether flying just above the surface or resting on it—be detected *before* or *within* the processing window defined by this reference. The key question, therefore, is not whether bats can detect prey over water, but how ensonification strategies are configured so that prey-induced deviations reliably emerge against a highly stable reference background.

Beam shape and ensonification geometry play a decisive role in resolving this paradox. Mouth-emitting bats such as *Myotis daubentonii* can regulate beam width through changes in gape and call frequency composition, thereby controlling the spatial extent and angular distribution of insonified space (***Siemers et al., 2001***; ***Schnitzler et al., 1994***). Crucially, field measurements show that *M. daubentonii* emits a markedly *narrower* sonar beam during water-surface foraging than would be expected from laboratory measurements or from aerial hawking behaviour (***Surlykke et al., 2008***). This narrow-beam strategy limits the amount of acoustic energy incident on the water surface at higher frequencies, thereby constraining the strength and variability of surface echoes while preserving forward sensitivity.

Over calm water, this frequency-dependent ensonification produces a highly structured echo scene, where the water surface provides a temporally stable reference dominated by lower-frequency components with broader beam widths. Higher-frequency components, emitted within a narrower forward beam, are preferentially reflected away at shallow incidence angles and therefore contribute less to the reference echo. Importantly, however, the effective temporal anchor for call timing need not correspond to the widest-angle or strongest surface reflection in a strictly geometric sense. As the sonar beam sweeps concentrically with minor fluctuations in head orientation, beam shape, and call structure, a range of near-surface scatterers—including weak surface ripples, capillary waves, or transient debris—may intermittently dominate the echo delay that governs inter-pulse timing.

Such a distributed anchoring mechanism offers a parsimonious explanation for the oscillatory structure in inter-pulse interval reported during water foraging (e.g. figure 5c, middle panel in Kalko and Schnitzler (1989b); figures 2–8, panels *d* in ***Schnitzler et al. (1994)***). Within the responsivity framework, these fluctuations reflect adaptive switching between closely spaced reference delays within a constrained near-field window. Crucially, this implies that the temporal anchor for call timing may arise from whichever echo source currently provides the most reliable and repeatable delay, rather than from a single fixed geometric feature such as the point at maximum/minimum beam incidence angle on the water surface.

**Figure 5.**
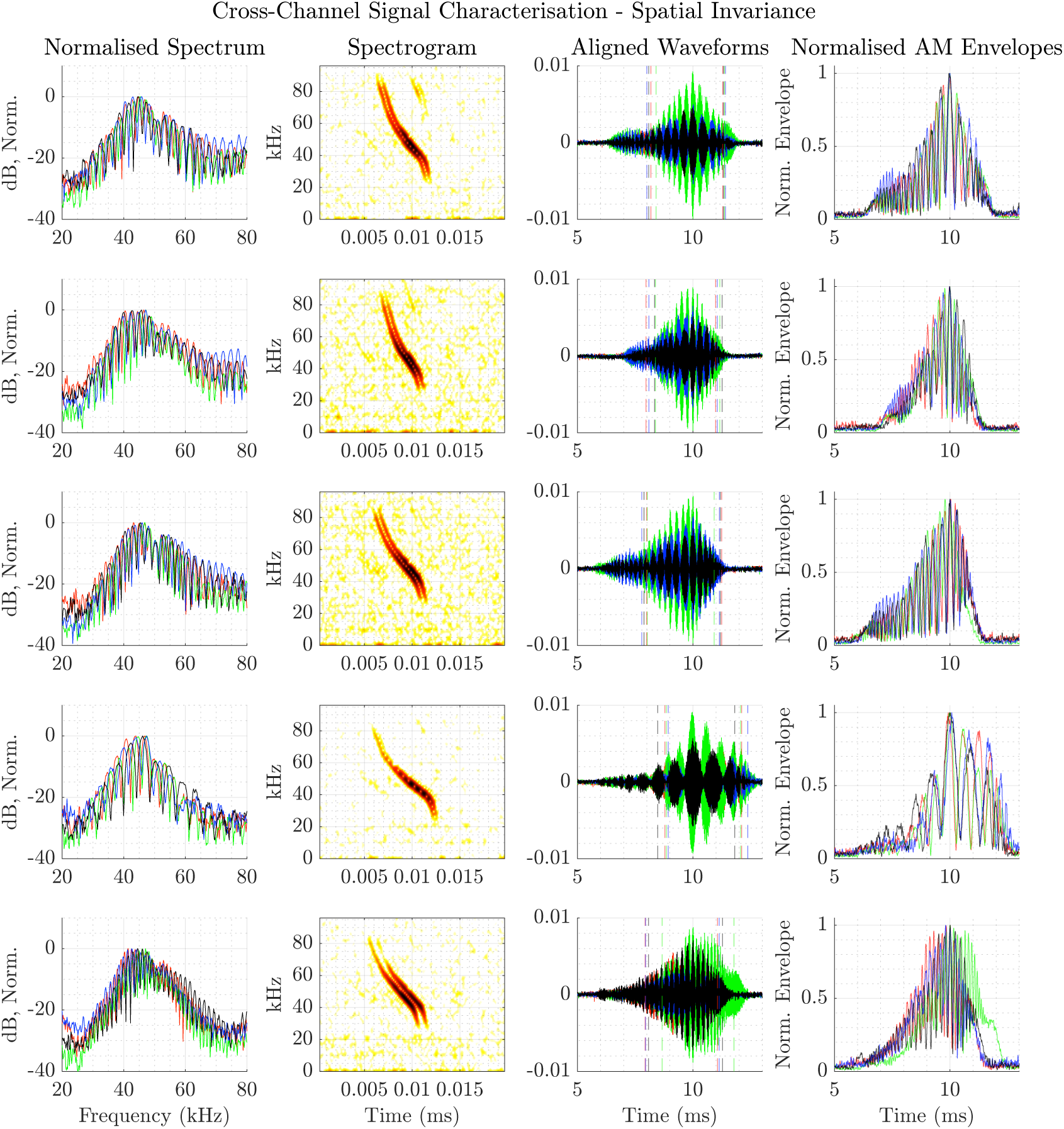
Cross-channel signal characterisation reveals spatial invariance of echo signals across array microphones. Each row corresponds to a call event recorded on a 4-channel microphone array, selected based on an amplitude threshold. Each line colour represents one of the four microphone channels (Red, Green, Blue, and Black corresponding to Channels 1–4). For each call, signals were aligned across channels using cross-correlation and centred on the peak energy. Left: Normalised power spectra from all four channels show a near-identical spectral shape. Centre-left: Spectrograms of the aligned reference channel show call time-frequency structure. Centre-right: Waveforms with energy-based duration markers (10–90% cumulative energy) show preserved temporal morphology. Right: Amplitude envelopes from all channels, normalised and low-pass filtered, confirm consistent envelope dynamics across spatial positions. These analyses confirm high spatial coherence and channel-invariant temporal-spectral structure of recorded signals, supporting the hypothesis of spatial invariance in ripple imprint.

This interpretation further emphasises that prey detection over water does not require prey echoes to replace the reference echo outright. Instead, prey-induced perturbations—whether preceding, coincident with, or embedded within the reference echo—can be detected as deviations from an expected temporal and spectral pattern. The precise nature of the reference point governing *T_*a*_* during foraging therefore remains an open question, and resolving it will require controlled experiments that independently manipulate surface structure, beam geometry, and echo delay. Nevertheless, the present framework provides a principled account of how stable call timing can coexist with a dynamically structured echo scene, and how oscillatory IPIs may naturally emerge from reference-based control rather than reflecting loss of prey tracking (See also related discussions of call grouping in ***Umadi and Firzlaff (2025)***).

Provided that the reference echo is sufficiently stable across successive calls, such perturbations are expected to stand out as violations of expectation rather than absolute intensity peaks. This interpretation aligns with evidence for object-specific adaptation and sensitivity to acoustic mismatch in the bat auditory system, whereby neural responses rapidly adapt to repeated echo patterns while remaining responsive to subtle, behaviourally relevant deviations (***Wiegrebe and Schmidt, 1996***; ***Sanderson and Simmons, 2000***; ***Pastyrik and Firzlaff, 2022***). Calm water maximises this predictability, whereas surface roughness introduces temporal and spectral variability that degrades the contrast between reference and prey echoes, consistent with the field observations by ***Schnitzler et al. (1994)*** in *N. leporinus*.

A flying insect hovering slightly above the water surface may return an echo that precedes the surface echo, whereas an insect resting on or contacting the surface will generate perturbations embedded within the reference echo but altered by body orientation, wing motion, or transient surface disturbance. ***Schnitzler et al. (1994)*** and related works (***Trappe and Schnitzler, 1982***; ***Von Der Emde and Schnitzler, 1990***; ***Schnitzler and Kalko, 2001***; ***Moss and Surlykke, 2001***) emphasise that prey motion must be an important clue to distinguish prey from inedible debris, with temporary glints and spectral cues distinct to live insects and fish breaking the water. On the other hand, water foraging *M. daubentonii* have been reportedly seen picking up leaves of a willow tree by Kalko and Schnitzler (1989a), and Schnitzler et al. (1994) report cases of discarded presumed prey, showing that the bat’s ability to *identify* the prey could involve active learning.

Field recordings analysed here are consistent with these considerations (figure 7). Across 54 call sequences, call rates and responsivity-inferred cue distances clustered around an effective reference distance of ∼2.5 m, corresponding to a substantial ensonification volume rather than to the bat’s instantaneous flight height above the water. This is notable given that *M. daubentonii* is typically reported to fly 30–40 cm above the surface during water-foraging (Kalko and Schnitzler, 1989a), a value far smaller than the distances inferred from call timing. As in the reanalysis of *N. leporinus*, where flight height decreases from ∼20 cm at 2 m range to 6–8 cm at 1 m without corresponding compression of echo–pulse delay (***Hartley et al., 1989***), the present field data indicate that flight height itself does not serve as the primary regulator of call rate over most of the foraging bout.

**Figure 6.**
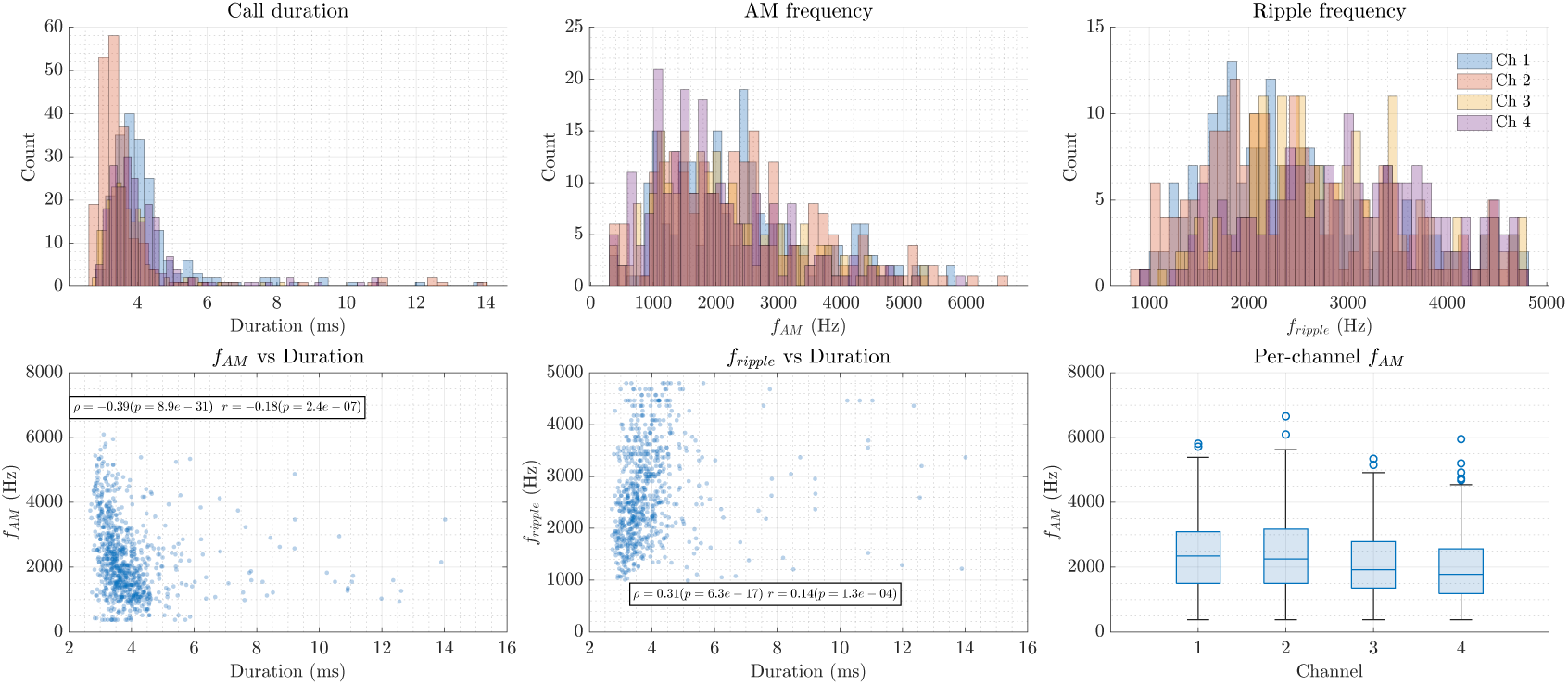
Summary of acoustic interference analysis across four recording channels (*N* = 207). Top row: histograms of call duration (per channel, transparent overlay), amplitude modulation (AM) frequency, and ripple frequency estimates. Bottom row: scatterplots showing the relationships between call duration and AM frequency (left), and between call duration and ripple frequency (middle). Right: per-channel boxplots of AM frequency distributions. Channel-specific histograms use partially transparent colours to highlight overlaps. Together, these plots illustrate that AM and ripple frequency estimates are relatively stable across channels, whereas call duration exhibits systematic variation, consistent with geometric differences in source–receiver configuration.

**Figure 7.**
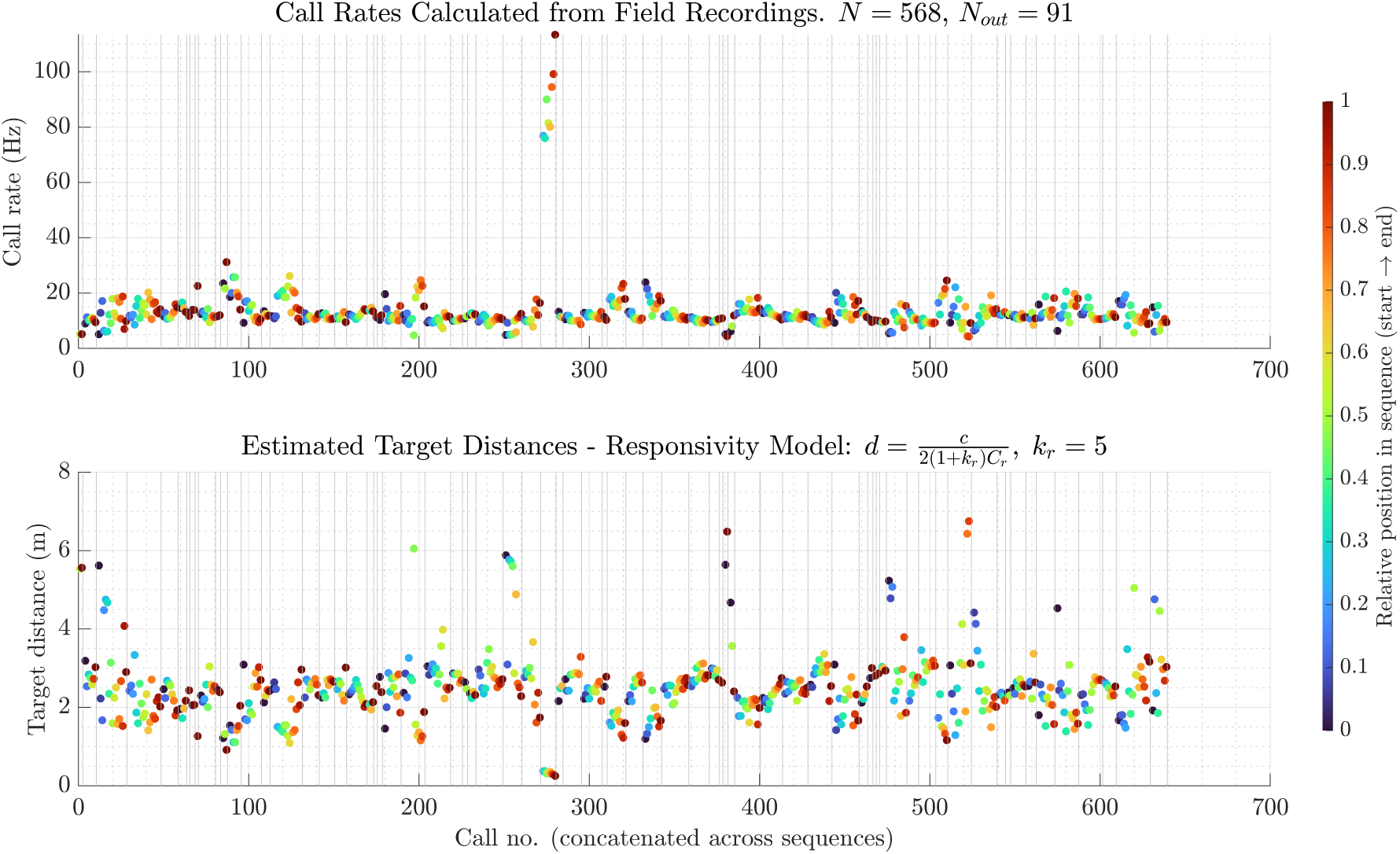
A total of 54 call sequences were analysed, comprising 568 validated calls and 91 excluded as outliers (d>10 m). A small proportion of the recordings contained calls from *Pipistrellus* individuals foraging near the array. Calls were manually extracted and validated for signal quality; in most sequences, the terminal buzz calls did not provide sufficient SNR to be included. Call rates were calculated only for continuous validated calls. The responsivity equations, when rearranged, provide a direct estimate of target distance (see also figure 9). Assuming a responsivity coefficient of *k*_*r*_ = 5, estimated target distances are shown in the lower panel. Most call rates, corresponding to the foraging phase, cluster around a mean of ∼ 13.5 Hz, which corresponds to a target distance of ∼ 2.5 m. One notable sequence exhibited call rates approaching 113 Hz (near call number 270), corresponding to an estimated target distance of ∼ 25 cm. In both panels, the colour coding indicates the relative position of a call within its sequence (from start to end), with thin grey vertical lines marking sequence boundaries. Each coloured point in the top (call rate) and bottom (distance) panels represents the same validated call. The results indicate that water-foraging *M. daubentonii* may fly over water bodies across a considerable range of heights. However, the target may not necessarily be the water surface itself; since these calls were recorded from the shore, the bats may also have used shorelines and surrounding vegetation as acoustic references in passing. When cross-referenced with the ripple-frequency and envelope-AM frequency distributions, the corroborated values provide a line of evidence that these bats likely forage well above the 0.4 m previously reported flight height.

Terminal buzzes were largely absent from the field dataset, so the analysed calls predominantly represent the search and early approach phases. Under the responsivity framework, this implies that the temporal anchor governing *T_*a*_* is located ahead of the low-flying bat, rather than being tied to the immediate water surface beneath it. Taken together with behavioural descriptions of water foraging in *N. leporinus* (***Schnitzler et al., 1994***), these results strongly suggest that call timing is regulated relative to a forward, scene-level reference—potentially arising from distributed near-surface scatterers—rather than being directly controlled by flight height.

Further, ***Rydell et al. (1999)*** showed that *M. daubentonii* preferentially forages over calm water and avoids even mildly turbulent surfaces, despite higher insect abundance in disturbed areas. Under the present interpretation, such avoidance need not reflect an inability to detect prey per se, but rather a breakdown of the stable reference required for reliable deviation-based detection. Habitat-use studies further report switching between water-surface and aerial foraging modes, particularly near shorelines or vegetation (***Todd and Williamson, 2019***; ***Todd and Waters, 2007***), where alternative persistent references can substitute for the water surface.

The echo-acoustic mirror effects discussed by ***Genzel et al. (2015)*** can be accommodated within this view, particularly for extended, stable objects near shorelines such as leaves or emergent vegetation. However, mirror images of insect prey are unlikely to play a dominant role in water-surface foraging, as prey are typically small, irregularly oriented, and transient. Accordingly, the present framework emphasises first-order, scene-level predictability: the water surface structures the echo scene. At the same time, prey detection arises from deviations within this structured field rather than from stable prey-generated mirror images.

Collectively, these considerations suggest that a coordinated strategy of ensonification control, reference-based timing, and neural sensitivity to deviation enables water-surface foraging. The narrow sonar beams documented in field conditions (***Surlykke et al., 2008***) appear to be a key behavioural adaptation that stabilises the reference echo while maintaining sensitivity to prey-related perturbations. The rarity of this foraging mode across bat taxa reflects the convergence of multiple constraints, including fine control of beam geometry, flight stability at shallow incidence angles, and the ability to detect subtle echo deviations without losing temporal control.

Reinterpreting Call-rate Control in ***Noctilio leporinus***

To evaluate how water-foraging bats regulate call timing in scenes containing multiple potential acoustic references, I reanalysed the prey-capture data reported by ***Hartley et al. (1989)*** within the responsivity framework. In their original study, echo–pulse delay was interpreted under the implicit assumption that bats continuously tracked the prey item (a suspended mealworm) as the dominant reference throughout the approach. My reanalysis revisits this assumption in light of a model that explicitly links inter-pulse interval (IPI) to echo delay via a scaling factor *k_*r*_*.

When the measured IPIs from Hartley et al. are converted into an implied cue distance using responsivity equations, the inferred distances consistently fall below the reported bat–prey distance for most of the approach phase (figure 9). This discrepancy indicates that, during large portions of the flight, call timing was likely governed by echoes arising from a nearer reference than the mealworm. Given the experimental geometry—a bat flying over a masonite surface in a confined and familiar environment—the most parsimonious interpretation is that near-field echoes from the surface or other stable environmental features dominated timing control, with prey-centred tracking becoming relevant only at very close range.

**Figure 8.**
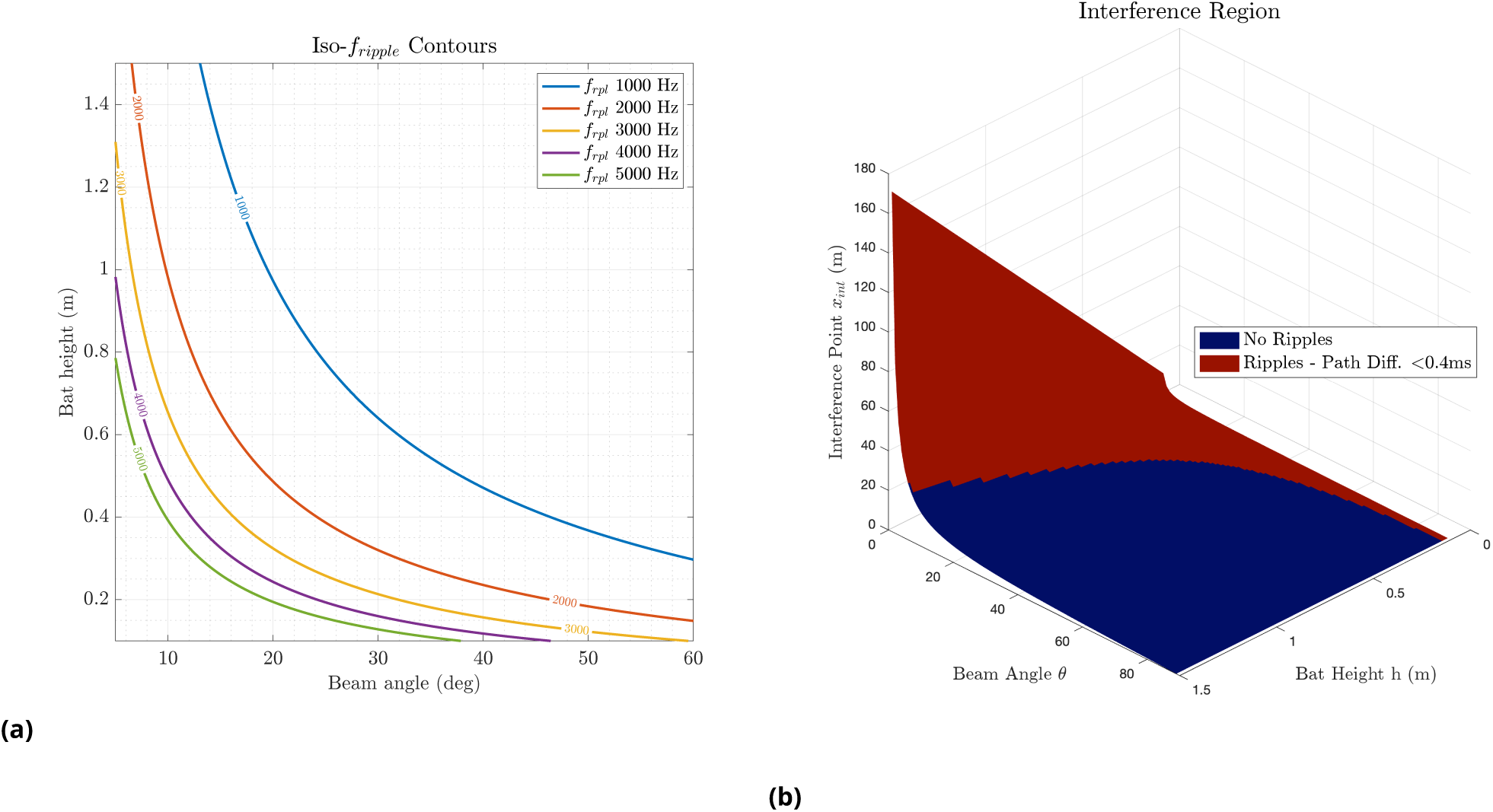
(a) Iso–ripple frequency contours predicted by the geometric interference model. Each coloured line indicates the locus of bat height–beam angle combinations that yield the same ripple frequency (1–5 kHz in 1 kHz steps). These contours illustrate the degeneracy of possible spatial configurations that can produce identical ripple frequencies, highlighting the need to consider additional acoustic cues (e.g. envelope shape, duration inflation) when inferring bat positions. (b) The plot shows the interference point location *x*_int_ as a function of bat height (*ℎ*) and beam angle (*θ*) assuming planar reflection and a fixed shore distance. The blue surface represents all possible configurations where the reflected beam intersects the line to the microphone, while the overlaid red region marks the subset of conditions where the delay between the direct and reflected paths is less than 0.4 ms.

**Figure 9.**
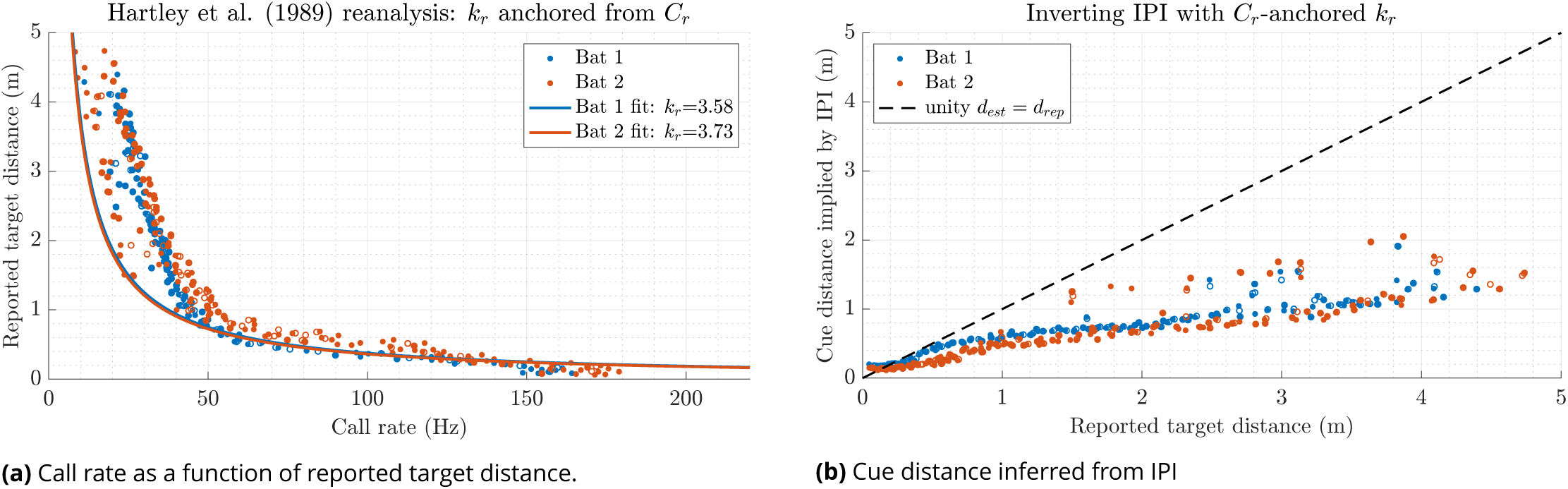
Reanalysis of call timing in ***Hartley et al. (1989)***. (a) Digitised call-rate data for two *N. leporinus* individuals plotted against reported bat–target distance. Solid curves show the responsivity model prediction using a responsivity coefficient *k*_*r*_ anchored from the high-rate, near-target regime of the call-rate data (see Methods). Although the terminal call rates are matched by construction, the model does not account for the full call-rate–distance scatter under the assumption of continuous prey-distance locking. (b) IPIs digitised independently from call-rate data, converted to an implied cue distance (equation (16), using the call-rate–anchored *k*_*r*_ values. The dashed line indicates equality between inferred cue distance and reported target distance. Filled symbols represent directly digitised measurements, whereas open symbols indicate distance-aligned interpolations introduced to form the union of independently digitised call-rate and IPI datasets. Points lying systematically below the unity line indicate that call timing was governed by echoes originating substantially closer than the reported prey distance over most of the approach, with convergence toward prey-centred tracking only at short range.

**Figure 10.**
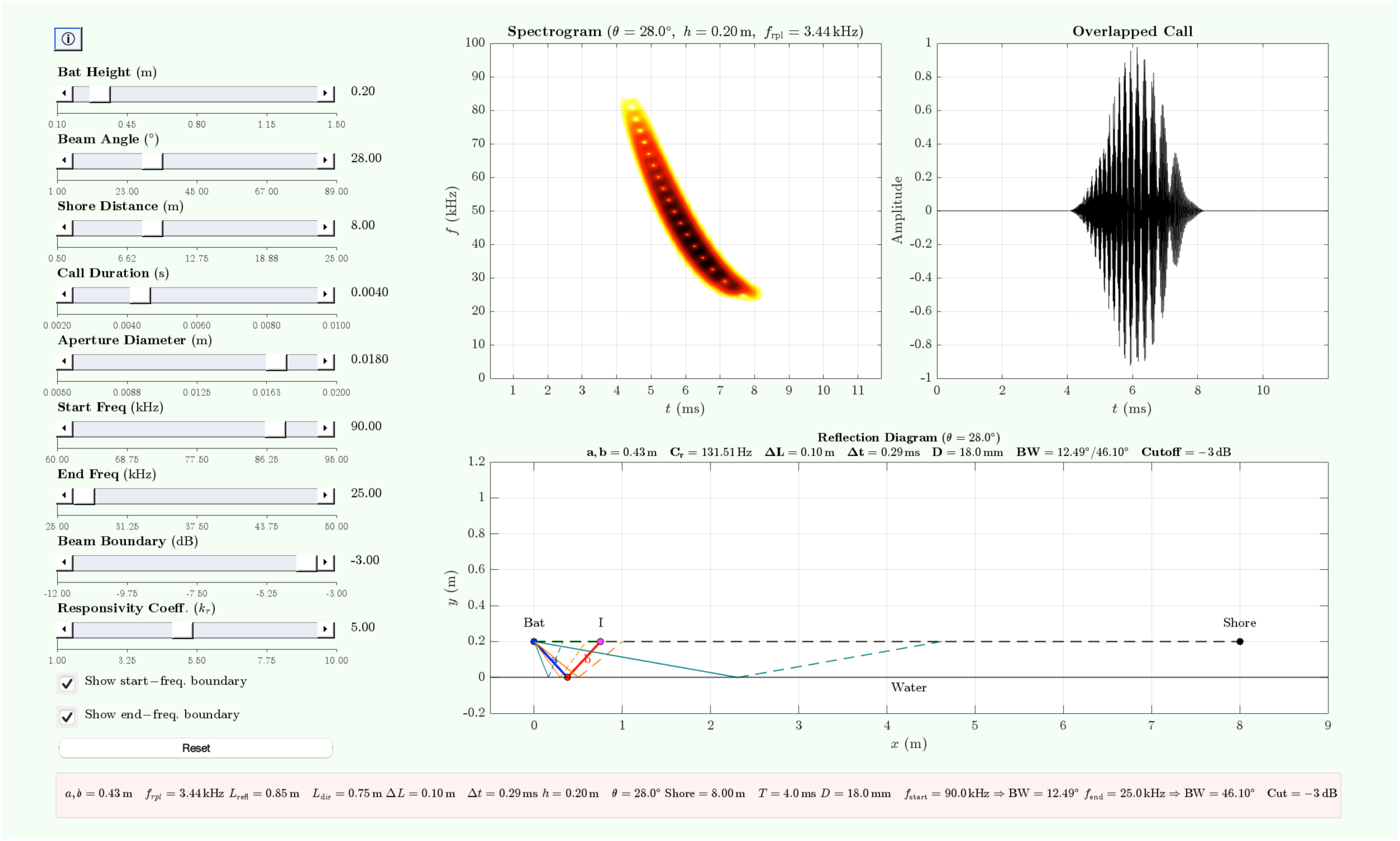
*Ripple Studio:* interactive app for modelling water-surface call–echo interference with responsivity-based timing control. The app simulates spectral interference in bat echolocation over water by overlapping a direct FM call with its specularly reflected copy from a flat surface. The overlapped signal is visualised as a spectrogram (top left), time waveform (top right), and a geometric reflection diagram (bottom), with interactive controls for call structure, emitter geometry, and responsivity. The model incorporates a responsivity-based call-timing framework: the effective call rate (*C*_*r*_) is computed from an inferred effective slant range (*d*_eff_) determined by the lateral beam footprint at the water surface and the bat’s height.

An alternative interpretation of the ***Hartley et al. (1989)*** data is that bats may indeed have tracked the prey continuously throughout the approach, but that the effective responsivity coefficient *k_*r*_* itself varied systematically with distance or behavioural phase. Under this view, the elevated call rates observed at larger bat–prey distances would not reflect the use of a nearer acoustic reference, but instead a dynamic adjustment of the mapping between echo delay and call timing. Such flexibility could arise from changes in sensory gain, attentional allocation, or processing latency as the task demands evolve during prey capture. In principle, a distance-dependent or context-dependent *k_*r*_* could reconcile the observed call-rate–distance relationship with strict prey locking.

However, the ***Hartley et al. (1989)*** dataset provides limited leverage for discriminating between reference switching and a distance-dependent *k_*r*_*. A model in which *k_*r*_* varies smoothly with target distance is difficult to constrain empirically without independent manipulation of reference distance or echo structure, and would require coordinated, systematic changes in processing latency that mirror prey distance throughout the entire approach, despite the presence of strong and stable near-surface echoes. By contrast, the reference-switching interpretation parsimoniously accounts for both the elevated call rates observed at larger reported distances and the late convergence of inferred cue distance toward the prey (figure 9), without invoking continuous recalibration of the sensorimotor mapping. Accordingly, while the data do not definitively exclude the possibility that *k_*r*_* is dynamically modulated, they are most consistent with a framework in which *k_*r*_* acts as a relatively stable scaling between echo delay and call timing, and behavioural transitions arise primarily through changes in the effective acoustic reference governing *T_*a*_*. Definitive tests of distance-dependent responsivity will require experiments that explicitly decouple prey distance from other acoustic references and manipulate echo delay independently of target geometry.

The structure further supports this interpretation in figure 16 of ***Hartley et al. (1989)***, which plots echo–pulse delay as a function of bat–target distance. At larger distances, the data show substantial scatter: echo–pulse delays do not collapse onto a single curve predicted by strict target locking. Such variability is expected if bats intermittently reference different echo sources with distinct delays, rather than maintaining continuous prey-centred control. Importantly, this scatter diminishes as the bat approaches the target, and at distances below approximately 0.5 m the echo–pulse delay flattens despite continued increases in call rate (figure 15, ***Hartley et al. (1989)***). Within the responsivity framework, this flattening is consistent with a regime in which *T_*a*_* becomes very small and the inter-pulse interval is increasingly dominated by processing and motor constraints rather than by further reductions in geometric delay.

***Hartley et al. (1989)*** computed echo–pulse delay as the time between echo reception and emission of the subsequent call, but did not explicitly incorporate call duration or echo duration into their timing model. This effectively treats the relevant echo as an instantaneous event, an assumption that is reasonable for their behavioural analysis but obscures how processing time scales with echo delay. In the responsivity formulation, the observed echo–pulse delay corresponds to (1 + *k_*r*_*)*T_*a*_*, such that the near-constant delays observed at close range imply a roughly constant multiplicative scaling between sensory delay and motor output. While this does not constitute a direct measurement of *k_*r*_*, it provides an empirical constraint: the data are difficult to reconcile with a model in which *k_*r*_* varies strongly with distance or behavioural phase.

Crucially, the flattening of echo–pulse delay at short range (figure 16, ***Hartley et al. (1989)***) suggests that once the prey echo becomes reliably dominant, further reductions in distance are accommodated primarily through increases in call rate, reduction of call duration (figures 6 & 7, ***Hartley et al. (1989)***) within a constrained processing window. This interpretation aligns with my reanalysis (figure 9a), where measured call rates exceed predictions based on prey distance alone but are well captured when a nearer reference is assumed for most of the approach. The late convergence of inferred cue distance toward the reported target distance thus marks a reference switch.

The experimental paradigm of ***Hartley et al. (1989)*** is therefore uniquely informative for the present study. By combining controlled prey presentation with measurements of bats’ position and rich echolocation call parameters, it provides rare access to the temporal logic of biosonar control in a water-foraging context. When viewed through the lens of responsivity, their data offer a coherent picture in which bats exploit stable environmental references to regulate call timing over much of the approach, reserving prey-centred tracking for the final interception phase. The apparent constancy of the scaling between echo delay and call timing across this transition is consistent with a fixed *k_*r*_*. However, confirming this interpretation will require experiments that explicitly manipulate reference distance and echo structure.

Physiological Implications of the Deduced Responsivity Coefficient *k*_*r*_ in ***Noctilio leporinus***

The responsivity estimates obtained here also have important physiological implications for call production in *Noctilio leporinus*. For the estimated value of *k_*r*_* ≈ 4 (see section), the responsivity model predicts a maximum call rate of ≈ 650–700 Hz at a presumed capture distance of *d* ≈ 5 cm (equation (2)). Such call rates have never been reported in bats, either in laboratory or field studies, including during the terminal buzz.

This discrepancy suggests that although the responsivity framework permits extremely high call rates at very short ranges, bats are unlikely to exploit this theoretical limit in natural behaviour. Instead, call rates appear to saturate at substantially larger distances (on the order of 20–30 cm), consistent with a regime in which further increases in repetition rate would produce excessive temporal overlap between calls and echoes or impose constraints on call production and auditory processing.

Physiological evidence supports this interpretation. Studies of laryngeal phyiology demonstrate that bats are capable of producing call rates approaching 160–200 Hz during the terminal buzz, but that these muscles operate near their mechanical limits at such rates (Suthers, 1988; ***Elemans et al., 2011***). Importantly, call durations at this stage are extremely short (typically ≲ 0.5–1 ms), restricting the temporal window in which further rate increases would remain functional. Field observations and controlled experiments likewise report species-specific terminal call rates that vary widely but remain far below the theoretical maxima implied by responsivity alone (Kalko and Schnitzler, 1989a; ***Hartley et al., 1989***; ***Schnitzler et al., 1994***).

Within the responsivity framework, this saturation emerges naturally: as distance decreases and echo delays approach the minimum processing window, further reductions in *T_*a*_* contribute little to shortening the inter-pulse interval. Instead, behavioural control transitions to a regime dominated by physiological and biomechanical constraints, rather than by geometric delay. Thus, the absence of ultra-high call rates reveals where the sensorimotor loop becomes capped by the limits of vocal production and echo segregation.

These considerations suggest that while *N. leporinus* may possess the physiological capacity for rapid call production, ecological and biomechanical factors ensure that this capacity is rarely, if ever, fully expressed. Responsivity, therefore, provides an upper bound on call timing that is modulated in practice by anatomy, echo overlap, and task demands, rather than a target that bats attempt to reach during prey capture.

### Geometric–Responsivity Inference of Foraging Behaviour

A central implication of the present analysis is that geometric models of echo formation, when combined with the responsivity framework, provide a powerful means of inferring foraging behaviour even in situations where direct spatial localisation of bats is impractical. In water-foraging bats, frequency-modulated calls recorded close to the water surface often exhibit pronounced spectral ripple patterns arising from interference between the direct call and its surface-reflected component. As demonstrated by iso-ripple analyses (figure 8), however, these spectral structures are inherently non-unique. Spectral ripples alone therefore do not uniquely encode bat height or orientation, and cannot be used in isolation to reconstruct flight geometry – a caution warranted from early studies of Kalko and Schnitzler (1989b).

This geometric ambiguity reflects a fundamental identifiability problem. Because the orientation of the bat and the acoustic axis of emission are typically unknown, and because the same interference delay can arise from multiple height–angle combinations, ripple frequency provides only a constraint on the effective path-length difference, not on the underlying spatial configuration. Additional signal features are therefore required to narrow the set of plausible solutions.

These include call intensity, spectral bandwidth, and the consistency of ripple structure across successive calls, which together can help eliminate geometries that are acoustically implausible or behaviourally unstable over time. In practice, directional microphones further restrict the recorded signal set by preferentially capturing calls emitted within a limited angular sector, reducing the contribution of low-amplitude or off-axis calls that would otherwise confound inference.

Within this context, the responsivity framework offers a complementary behavioural constraint. By linking call timing directly to echo delay via a scaling factor *k_*r*_*, responsivity imposes temporal structure on otherwise ambiguous geometric interpretations. While multiple height–angle combinations may produce the same instantaneous ripple pattern, only a subset of these configurations will evolve consistently with the observed call-rate dynamics over a sequence of calls. Continuous call sequences, therefore, provide additional information: plausible trajectories must not only reproduce the observed ripple structure at individual time points, but also yield a temporally coherent evolution of echo delay and call timing. In this sense, responsivity acts as a behavioural regulariser, allowing the most likely geometric interpretations to be identified without requiring explicit localisation.

This approach is particularly valuable because direct localisation of water-foraging bats remains challenging. As noted by ***Surlykke et al. (2008)***, strong surface reflections, low flight heights, and rapidly changing geometry degrade the performance of conventional microphone-array localisation methods. Recent methodological work has further shown that relative motion, beam dynamics, and multipath propagation can substantially bias localisation estimates even in dense arrays (Umadi, 2025f). Although sophisticated algorithms can partially mitigate these effects, they remain technically demanding and are not routinely accessible to field ecologists.

By contrast, behavioural inference based on geometric signal features and responsivity relies primarily on single-channel recordings or modest multichannel setups, making it substantially easier to deploy in the field. Single microphones are widely available and already form the backbone of many bat-monitoring programmes. At the same time, recent advances in low-cost, open-source multichannel ultrasonic hardware further lower the barrier to acquiring richer datasets (Umadi, 2025c,d). Importantly, the responsivity-based approach is internally self-consistent: it does not require absolute spatial reconstruction, but instead tests whether observed call timing and spectral structure are compatible with specific classes of behavioural strategies.

Additionally, as the water-foraging bats typically operate at relatively stable call rates, flying at stable heights over extended search-approach phases, such a behavioural context provides a natural testbed for evaluating predictions of wingbeat–call synchrony derived from temporal feasibility constraints. Linking the present responsivity–geometry framework with the temporal feasibility analysis (Umadi, 2025e) enables independent assessment of respiratory–locomotor coordination under well-defined sensory control regimes.

Thus, the combination of acoustic geometry modelling and responsivity provides a practical and conceptually grounded alternative for inferring foraging behaviour in complex acoustic scenes. This framework leverages invariant relationships between geometry, echo delay, and call timing to constrain behavioural interpretation. In doing so, it enables robust inference from data that are already routinely collected, and offers a tractable pathway for extending behavioural analyses of echolocation into challenging natural environments.

## Acknowledgements

I gratefully acknowledge the support and insightful guidance of my doctoral supervisor, Uwe Firzlaff. In particular, his early comments on the spectral ripple patterns observed in field recordings were instrumental in motivating and shaping the conceptual direction of this work. I also thank Diana Marcela Ochoa Sanz at *Instituto de Ecología A.C., Xalapa, México* for generously sharing her field observations and behavioural insights on *Noctilio leporinus*, which greatly informed the ecological interpretation of the results. I am further grateful to my colleagues for their valuable discussions, encouragement, and feedback throughout the development of the study.

## Ethics Statement

Field recordings were conducted in accordance with local regulations – no special permit was required. No animals were captured, handled, or experimentally manipulated during this study.

## Data and Code Availability

The simulation code, app, analyses scripts, and data are available via:

1. GitHub Repo: https://github.com/raviumadi/ripples
2. Zenodo Archive: https://doi.org/10.5281/zenodo.18207549

## Appendix 1

### Specular-reflection Accounted Geometry

See main text section **Methods: Modelling Spectral Interference in Water-Foraging Bats**

**Appendix 1—figure 1.**
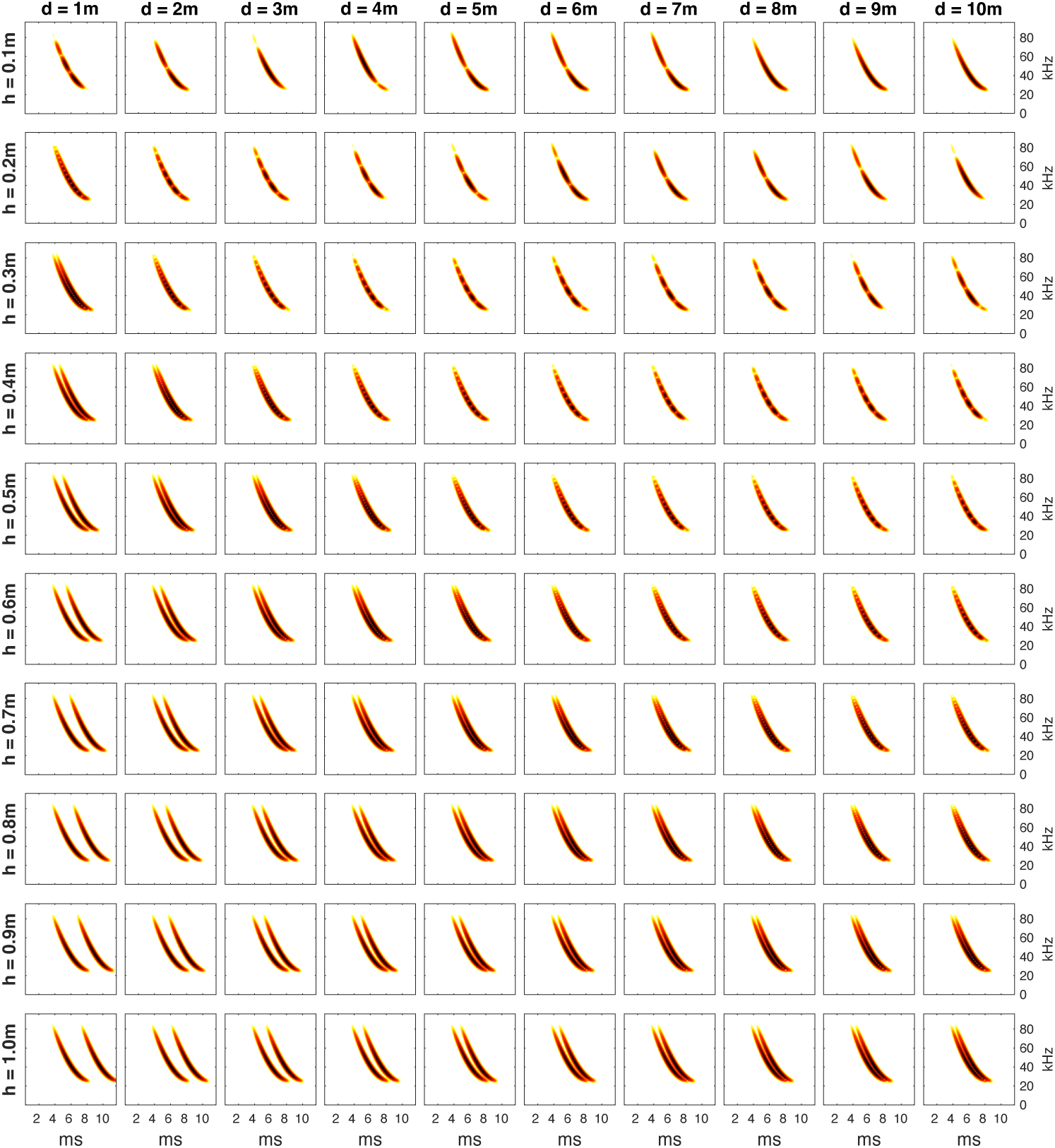
Specular reflection–accounted geometric model of call–echo interference across bat height and horizontal distance. Each panel shows the simulated spectrogram of a broadband FM echolocation call overlapped with its specularly reflected copy from a flat water surface, assuming mirror-symmetric reflection geometry. Rows correspond to bat height above the surface (*ℎ* = 0.1–1.0 m), and columns to horizontal distance (*d* = 1–10 m), which parameterises the geometric path difference between direct and reflected propagation. The temporal offset between the two paths is given by 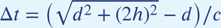 and determines the resulting spectral ripple spacing. For a given Δ*t*, the spectral ripple structure remains invariant with respect to receiver position, indicating that beyond a given distance the interference pattern is set by geometry rather than by receiver location. Varying *d* in this model, therefore, implicitly alters the angle of incidence and effective beam–surface interaction, rather than modelling receiver-dependent effects. Because identical values of Δ*t* can arise from multiple combinations of *ℎ* and *d*, the resulting ripple structure is geometrically degenerate with respect to flight height and beam incidence angle (an *isoripple* condition). Consequently, spectral ripple patterns alone do not uniquely specify bat height, orientation, or receiver position, even when specular reflection geometry is explicitly accounted for.

## Appendix 2

### At-Receiver Path Difference Geometry

See main text **section Methods: Contrasting Geometric Model Without Reflection Symmetry**

**Appendix 2—figure 1.**
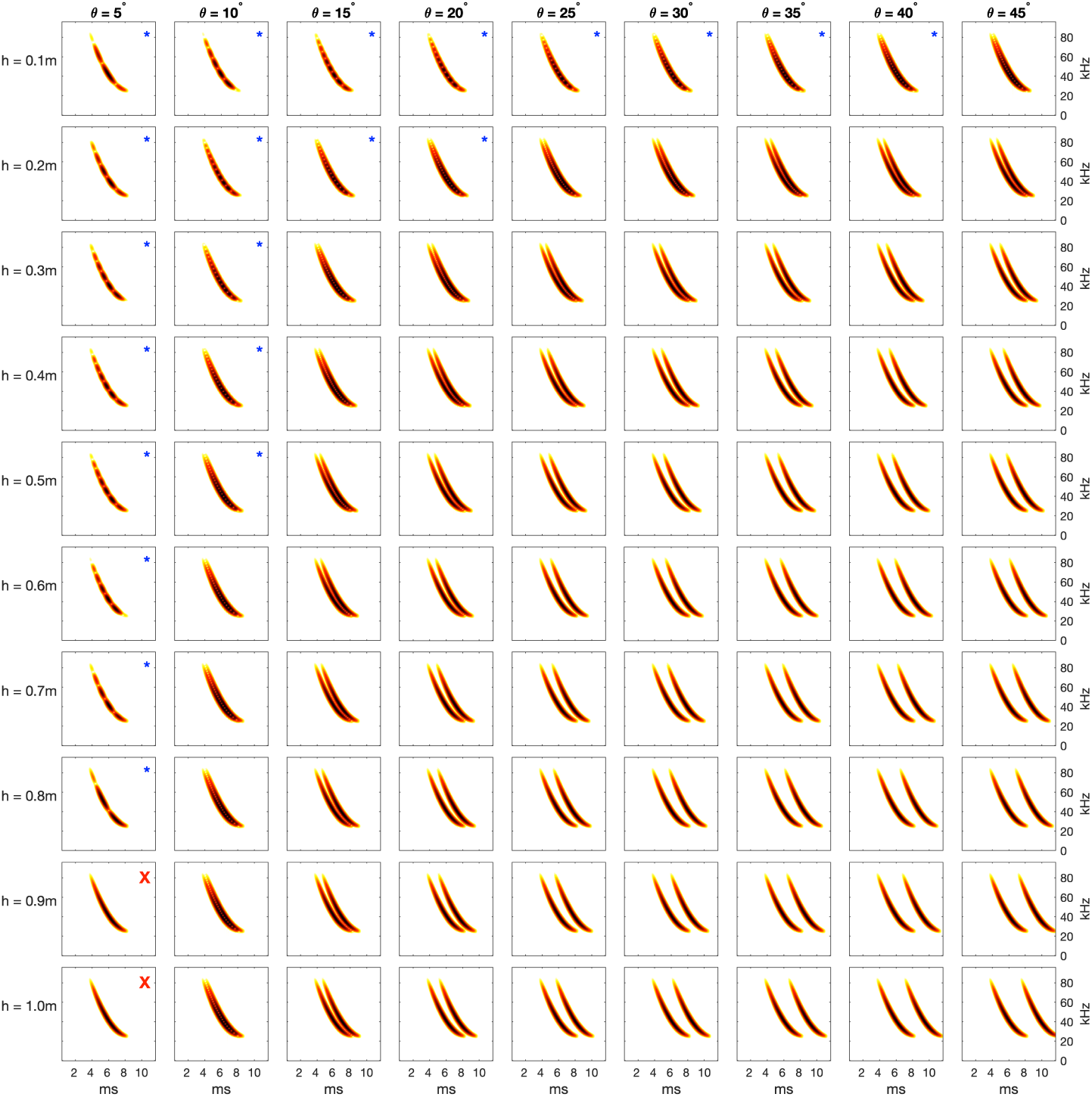
Evaluation of the at-receiver interference model for spectral ripple formation under varying flight geometries. Simulated overlapped echolocation calls are shown for combinations of bat height above the water surface and horizontal distance to the receiver, following the at-receiver interference formulation used in earlier studies (e.g. *Kalko and Schnitzler* (*1989b*); *Ratcliffe and Jakobsen* (*2018*)). In this model, spectral ripples arise from interference between a direct call and a surface-reflected copy arriving at the receiver with a fixed path-length difference determined solely by height and range, without explicit consideration of emission angle or reflection geometry. Ripple structure emerges only within a restricted region of parameter space corresponding to low bat heights and shallow effective incidence angles (blue asterisks), where the assumed path difference is approximately consistent with specular reflection. Outside this regime, interference patterns weaken or disappear, indicating that the model does not produce physically plausible reflections for many height–distance combinations. This figure illustrates that the at-receiver interference model is valid only under constrained geometric conditions and cannot, by itself, support general inference of flight height or beam orientation. These limitations motivate the use of explicit geometric reflection models combined with call-timing constraints, as developed in the main text.

## Appendix 3

### *Ripple Studio* User Guide (v1.2)

*Ripple Studio* is an interactive MATLAB app designed to demonstrate and quantify spectral interference (“ripples”) produced when a bat echolocation call propagates above a smooth, specularly reflecting water surface. The app implements a forward model of (i) two-path geometry (direct and surface-reflected propagation), (ii) frequency-dependent beam spreading under a circular-piston approximation, and (iii) time-domain overlap of direct and delayed copies of a down-swept FM call. Ripple Studio provides three linked outputs that update in real-time as parameters are adjusted: a spectrogram of the overlapped signal, the corresponding time waveform, and a plan-view geometry diagram, including optional beam-boundary rays.

*Ripple Studio* is intended both as (1) an explanatory tool for understanding the physical origin of spectral ripples and (2) a practical analysis aid for interpreting field recordings, because ripple spacing provides a deterministic signature of the underlying path delay.

### Installation and launching

*Ripple Studio* is implemented as a MATLAB class. Standalone installation packages are provided for macOS and Windows. MATLAB Runtime (freeware) is installed along with the packages for operation. The programmatic implementation exposes the function variables for advanced users.

#### Requirements

MATLAB R2018b or later is recommended; MATLAB R2021a or later is preferred for robust figure exporting with exportgraphics. No additional toolboxes are required beyond standard MATLAB signal and plotting functions.

#### Launching the GUI

From the MATLAB command window, ensure RippleStudio.m is on the path and run:

~~~
h = RippleStudio; % launches the GUI
~~~

The GUI opens as a fixed-size figure window containing the control panel (left) and three axes (right).

### Interface layout

#### Axes

The GUI contains three axes:

- **Spectrogram** (h.ax1, top-left): magnitude STFT of the overlapped signal, displayed in dB with a fixed dynamic range.
- **Waveform** (h.ax1b, top-right): time-domain overlapped signal.
- **Geometry diagram** (h.ax2, bottom): direct and reflected paths, the reflection point, the interference point, and optional beam-boundary rays.

#### Controls

The left panel contains sliders (with numeric readouts), two checkboxes controlling beam-boundary overlays, a **Reset** button, and a help menu (**Help → Show description…**) plus a small “i” button.

### Controls and what they mean

All slider adjustments trigger an immediate redraw of all three axes. The current parameter set and derived quantities are also stored in h.param (Section S7).

#### Geometry sliders Bat Height (m)

Emitter altitude above the water surface. Increasing height increases the typical path-length difference between direct and reflected propagation for a fixed beam angle, thereby changing the interference delay *τ* and ripple spacing *f*_ripple_.

#### Beam Angle (^◦^)

Downward aiming angle of the sonar axis relative to the horizontal. This controls the specular reflection point on the surface and therefore the direct and reflected path lengths. Shallow angles (slight downward tilt) typically push the reflection point farther away; steeper angles bring it closer.

#### Shore Distance (m)

A geometric reference distance used only for the diagram (horizontal extent and placement of a “shore” marker). It does not change the interference physics unless it constrains how you visually interpret the scene.

#### Call and source sliders Call Duration (s)

Duration of the synthetic down-swept FM call. This affects the temporal overlap of direct and delayed copies and the visibility of ripples in the spectrogram: ripples are most salient when the delay *τ* falls within the call duration such that the two copies overlap.

#### Aperture Diameter (m)

Effective emitter size used in a circular-piston model to compute frequency-dependent beamwidth. Larger apertures produce narrower beams (smaller beamwidth). Beamwidth is calculated separately at the start and end frequencies and used for visual boundary rays and an “effective” distance estimate used in the displayed call-rate value.

#### Start Freq (kHz) and End Freq (kHz)

Start and end frequencies of the down-swept FM call. These influence (i) the call’s time– frequency trajectory, (ii) beamwidth via the piston approximation (shorter wavelengths narrow the beam), and (iii) how ripple structure appears across frequency bands.

#### Beam Boundary (dB)

Amplitude cutoff used to define beam edges (e.g., −3 dB). More negative values widen the boundary rays. Beam boundary is used only for beam visualisation and beamwidth reporting (it does not change two-path delay *τ* directly).

### Responsivity slider

#### Responsivity Coeff. (*k*_*r*_)

Scaling coefficient that links inter-pulse interval (IPI) to echo delay *T_*a*_* under the responsivity framework:

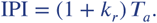

*Ripple Studio* reports an effective call-rate value *C_*r*_* computed from an effective distance estimate *d*_eff_ and the responsivity relationship

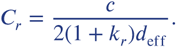

In the current implementation, *d*_eff_ is derived from the geometry and beam footprint (as displayed in the geometry panel). This value is intended as an interpretable proxy for how geometry and beamwidth jointly affect timing predictions.

### Beam-boundary checkboxes

Show start-freq. boundary / Show end-freq. boundary.

These overlays draw beam boundary rays (and their specular reflections) at the start and/or end frequency beamwidths, using the selected dB boundary. They help visualise how frequency-dependent directivity changes the ensonified footprint on the water surface.

### Reset and help

#### Reset

Restores all sliders to their default values and redraws the plots.

#### Help menu and info button

Opens a scrollable documentation window. If a documentation.txt file is present in the same directory as RippleStudio.m, its contents are displayed. Otherwise, a placeholder message is shown.

### Interpreting the plots

Spectrogram (overlapped call).

The spectrogram shows the magnitude of the short-time Fourier transform of the overlapped signal

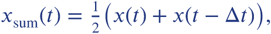

where *τ* is the geometric delay. The title reports *θ*, ℎ, and the predicted ripple spacing

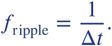

In the frequency domain, ripple null-to-null spacing is approximately *f*_ripple_, whereas in the time domain the beat frequency corresponds to

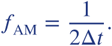

Ripple visibility depends on overlap: when *τ* is small relative to call duration, ripples become prominent; if *τ* is large, the two copies separate in time and interference diminishes.

#### Waveform (overlapped signal)

The waveform panel plots *x*_sum_(*t*) directly. When overlap is strong, characteristic beating (envelope modulation) is visible. This provides an intuitive time-domain complement to the spectrogram: ripple spacing in frequency corresponds to a beat period in time.

#### Geometry diagram

The geometry diagram shows:

- The bat location, the reflection point on the water surface, and the interference point.
- The direct path from bat to the interference point.
- The reflected path decomposed into two segments (bat → reflection point, reflection point → interference point).
- Optional beam-boundary rays (start and/or end frequency), including their specular reflections.

The diagram title and bottom information box report key derived quantities, including segment lengths, total reflected and direct path lengths, Δ*L*, Δ*t*, beamwidths at start/end frequency, and the selected dB cutoff.

### Exporting figures

*Ripple Studio* supports exporting axes “as seen” (titles, labels, ticks, and formatting preserved).

### Recommended export function

Use the public method exportAxesExact:

~~~
h.exportAxesExact(h.ax1, ’spectrogram.pdf’); % spectrogram
h.exportAxesExact(h.ax1b, ’waveform.pdf’); % waveform
h.exportAxesExact(h.ax2, ’geometry.pdf’); % diagram
~~~

Optional name–value arguments can be passed to exportgraphics, e.g.:

~~~
h.exportAxesExact(h.ax1, ’spectrogram.pdf’, ’Resolution’, 600);
h.exportAxesExact(h.ax1b,’waveform.png’, ’Resolution’, 600);
~~~

For PDF export, the method defaults to image-style export to preserve the exact on-screen appearance.

### Notes on reproducible exporting

For manuscript figures, the most reproducible workflow is:

1. Set all sliders to the desired values (record h.param).

2. Export each panel using exportAxesExact.

3. Archive the exported files alongside a saved .mat file containing h.param.

### Access to signals and parameters

*Ripple Studio* is designed to be transparent: all generated signals and derived values are accessible programmatically from the object handle.

### Generated signals

After any update, the following public read-only properties contain the most recent signals:

- h.call: direct FM call (column vector).
- h.indirectCall: delayed/reflected copy (column vector).
- h.overlapCall: overlapped signal 0.5(call + indirectCall).
- h.t_ms: time vector in ms.
- h.dt_s: geometric delay Δ*t* in seconds.
- Example:

x = h.call;

x_ref = h.indirectCall;

x_sum = h.overlapCall;

t_ms = h.t_ms;

tau_s = h.dt_s;

### Parameter structure (h.param)

All current slider values and derived quantities are stored in the struct h.param. This includes (non-exhaustive): height, beam angle, call duration, aperture, start/end frequency, cutoff, *k_*r*_*, segment lengths (*a*, *b*), direct/reflected path lengths, Δ*L*, Δ*t*, ripple spacing *f*_ripple_, beamwidths at start/end frequency, and the reported call-rate estimate. Example:

p = h.param;

disp(p.theta_deg);

disp(p.f_rpl_kHz);

### Saving and reloading a configuration

To archive a configuration for reproducibility:

p = h.param;

save(’RippleStudio_config.mat’,’p’);

Later, reload and apply:

load(’RippleStudio_config.mat’,’p’);

h.setParams(p); % updates sliders and redraws GUI

### Programmatic control (batch use)

*Ripple Studio* can be controlled without manual slider adjustment using setParams, enabling scripted sweeps (e.g., exploring ripple spacing across heights or angles) while still using the GUI as a live visualiser.

Example: set geometry and call parameters.

~~~
h.setParams(struct(’h’,0.35,’theta_deg’,25,’call_duration’,0.004, …
’f_start_kHz’,90,’f_end_kHz’,25,’aperture’,0.018, …
’cutoff_db’,-3,’kr’,5));
~~~

### Example: parameter sweep with exported panels

~~~
for H = [0.15 0.25 0.35 0.45]
h.setParams(struct(’h’,H));
h.exportAxesExact(h.ax1, sprintf(’spec_h%0.2f.pdf’,H));
end
~~~

### Practical workflow for interpreting field recordings

A recommended workflow for using RippleStudio as an interpretive tool is:

1. Measure ripple spacing from a recording (e.g., spacing of spectral nulls or peaks) to obtain *f*_ripple_.
2. Use the model relationship *f*_ripple_ to constrain Δ*t*.
3. Adjust height and beam angle in *Ripple Studio* to identify geometries consistent with the observed ripple spacing, while checking whether the resulting overlap regime matches the observed time-domain structure.
4. Archive the inferred configuration by saving h.param and exporting the panels used for the inference.

### Notes, limitations, and interpretation

- *Ripple Studio* implements a specular two-path model with a smooth reflecting plane and does not simulate rough-surface scattering or clutter statistics.
- The displayed ripple spacing and delay derive from the geometric path-length difference and therefore represent the deterministic component expected under calm-water conditions.Beamwidth is computed using a circular-piston approximation; thus, aperture diameter is an effective parameter capturing emission directivity rather than a direct anatomical measurement.
- The responsivity-based call-rate value is presented as an interpretable proxy linking geometry and timing predictions; interpretation should follow the definitions and assumptions described in the manuscript.

